# Enhancing Recovery from Gut Microbiome Dysbiosis and Alleviating DSS-Induced Colitis in Mice with a Consortium of Rare Short-Chain Fatty Acid-Producing Bacteria

**DOI:** 10.1101/2023.09.11.556543

**Authors:** Achuthan Ambat, Linto Antony, Abhijit Maji, Sudeep Ghimire, Samara Mattiello, Purna C. Kashyap, Sunil More, Vanessa Sebastian, Joy Scaria

## Abstract

The human gut microbiota is a complex community comprising hundreds of species, with a few present in high abundance and the vast majority in low abundance. The biological functions and effects of these low-abundant species on their hosts are not yet fully understood. In this study, we assembled a bacterial consortium (SC-4) consisting of *B. paravirosa, C. comes, M. indica*, and *A. butyriciproducens*, which are low-abundant, short-chain fatty acid (SCFA)-producing bacteria isolated from healthy human gut, and tested its effect on host health using germfree and human microbiota-associated colitis mouse models. The selection also favored these four bacteria being reduced in abundance in either Ulcerative Colitis (UC) or Crohn’s disease (CD) metagenome samples. Our findings demonstrate that SC-4 can colonize germ-free (GF) mice, increasing mucin thickness by activating MUC-1 and MUC-2 genes, thereby protecting GF mice from Dextran Sodium Sulfate (DSS)-induced colitis. Moreover, SC-4 aided in the recovery of human microbiota-associated mice from DSS-induced colitis, and intriguingly, its administration enhanced the alpha diversity of the gut microbiome, shifting the community composition closer to control levels. The results showed enhanced phenotypes across all measures when the mice were supplemented with Inulin as a dietary fiber source alongside SC-4 administration. We also showed a functional redundancy existing in the gut microbiome, resulting in the low abundant SCFA producers acting as a form of insurance, which in turn accelerates recovery from the dysbiotic state upon the administration of SC-4. SC-4 colonization also upregulated iNOS gene expression, further supporting its ability to produce an increasing number of goblet cells. Collectively, our results provide evidence that low-abundant SCFA-producing species in the gut may offer a novel therapeutic approach to IBD.

## Introduction

In biological ecosystems, species composition typically follows a skewed pattern, wherein a small subset of species is highly abundant, whereas the majority are low in abundance or relatively rare. This phenomenon, first observed by Darwin in his groundbreaking work[1], is a common characteristic across diverse biological classes and geographies. The same skewed distribution was later recognized by Preston in bird and moth populations during the 20th century[2]. This skewed biodiversity distribution is also reflected in the composition of the human gut microbiota. An analysis of the gut microbiome composition of individuals across various continents revealed a bimodal species distribution[3]. In this distribution, a handful of highly abundant species exist on the left of the log-normal distribution, whereas a large number of rare species congregate on the right. Upon further exploration of species distribution across thousands of individuals, Lawley et al. discovered that the human gut microbiota is primarily dominated by approximately 20 species, with the remaining species present in lower abundances[4].

Therefore, the focus of human gut microbiome research has been primarily directed towards understanding the roles and implications of these dominant species. Numerous studies have found that these highly abundant species often play vital roles in gut functionality. One such species is *Bacteroides thetaiotaomicron*, which is a common inhabitant of the human gut. Extensive research on this organism has provided valuable insights into how dominant members of the microbiome contribute to maintaining health and well-being. *B. thetaiotaomicron* plays a key role in nutrient utilization in the gut, where it helps metabolize polysaccharides that are otherwise inaccessible to the host[5]. Another well-understood example of a highly abundant gut species is *Faecalibacterium prausnitzii* which has been recognized as one of the main butyrate producers in the intestine[6]. The presence of highly abundant species can also be problematic. A well known example for that is *Escherichia coli* which depending on the strain type could proliferate in the dysbiotic gut can cause mild to fulminant diarrhea[7].

Metagenomic investigations of the gut microbiome have revealed several thousand species spanning diverse populations, with the majority being identified as low-abundance species. Intriguingly, most of these rare species have not been cultured, and even among them, comprehensive functional and mechanistic studies are sparse. Despite possessing a reasonable understanding of certain dominant species residing in the gut, our knowledge of the role of these rare species remains largely underexplored. However, emerging research indicates that these low-abundance species may profoundly influence host health and serve as potent activators of gut immunity among other potential effects. Eventually, these species contribute to the total richness and diversity of the system. In contrast, loss of diversity has been associated with various enteric and systemic diseases [8], further suggesting that these species might play a significant role in protection against these diseases.

In non-microbial communities, higher rates of extinction have motivated interest in diversity-stability relationships [9–12]. The insurance hypothesis states that high species richness reduces the temporal variability of a given community property by insuring the community against various perturbations [13]. The same concept has been observed in microbial communities. For example, Lieven et al. showed in experimental microcosms that the degree of richness and evenness in a community prior to perturbation affects the subsequent response [14]. Later, the same was explained for conditionally rare taxa (CRT) in the gut microbiome [15].

The objective of our study was to delve deeper into the functional roles of rare species in the gut, particularly those capable of producing short-chain fatty acids (SCFAs). SCFAs were selected as the focal point because of their significant contributions to immune cell proliferation[16], differentiation[17], apoptosis[18], gut barrier integrity[19], gut motility[20], and host metabolism [21]. Among the characterized SCFAs, butyrate has demonstrated protective effects against immune disorders such as Intestinal Bowel Disorders (IBD) [22]. Although the direct administration of butyrate compounds as a therapeutic intervention against IBD has encountered obstacles related to delivery, exposure duration, and patient compliance, there is continued interest in the beneficial properties of butyrate[23, 24]. The success of using prebiotics for IBD treatment depends on the presence of butyrate-producing bacteria[25]. For instance, treatment with butyrate-producing *F. praunitzi* was found to restore the aberrant microbiota community in an IBD mouse model, whereas *Butyricicoccus pullicaecorum* attenuated colitis in rats[26]. Given the scarcity of functional studies on rare gut species, we employed germ-free (GF) and humanized mouse models to explore the interspecies interactions of a manageable consortium of four SCFA-producing rare species in the human gut. We reasoned the selection of the strains based on the lower abundance in healthy metagenome, presence in the previously formulated culture library, ability to produce SCFA specifically butyrate and its differential abundance in IBD metagenome samples. We also utilized dietary fibers as supplements in humanized mice experiment to increase the abundance of these SCFA-producing bacterial members, thereby expecting to see an improvement in disease outcomes. Our findings indicated that the rare species mix we utilized colonized GF mice and alleviated DSS-induced colitis. In humanized mice, the same mix expedited the recovery of microbiome beta diversity and enhanced host health following Dextran Sodium Sulfate (DSS) perturbation. Our study also demonstrates the existence of functional redundancy within the gut microbiome, where low-abundant short-chain fatty acid (SCFA) producers playing a crucial role as a resilience factor. This redundancy facilitates a faster recovery from dysbiotic states upon the administration of SC-4. The colonization by SC-4 not only accelerates recovery but also upregulates the expression of the iNOS gene, contributing to an increase in mucin producing goblet cells. Taken together, our findings suggest that enhancing the population of low-abundant SCFA-producing bacteria in the gut may represent a new therapeutic strategy for managing IBD.

## Results

### Selection of strains for Short Chain fatty acid producing consortium (SC-4)

In our previous study, we developed a microbiota culture collection from healthy human donors that accounted for over 70% of the functional capacity of the healthy gut microbiome[27]. This collection included both high and rare abundant species. To curate a collection of short-chain fatty acid (SCFA)-producing strains from our library, we focused on several criteria: their ability to produce SCFAs, their low abundance nature, and their differential abundance in disease datasets. Given the association of butyrate with inflammatory bowel disease (IBD), our selection was particularly guided by the strains’ capacity to produce butyrate. This determination was based on both experimental inference and existing literature. Recognizing that the genes responsible for butyrate production in bacteria are well-documented, we primarily used the presence of these genes as the criterion for species selection in our study.

First, we determined the percentage abundance of already reported butyrate-producing bacteria in the donor fecal samples used for developing our culture collection[27] **(Fig. 1A)** as well as in a publicly available healthy human shotgun metagenomics dataset[28] **(Fig. 1B)** [27, 29–32]. Consistent with prior reports, our analysis revealed that the well-known butyrate-producing species *F. praunitzi* was highly abundant in donor metagenomes. Nevertheless, the mean abundance of most putative butyrate-producing species was less than 0.1%. To constitute a manageable mix of strains for further study, we randomly selected four species with abundances below this threshold **(Supplementary Fig. 1A)**. The abundance of these selected strains, measured from the metagenome sequence data of six donor fecal samples, showed that all four bacteria exhibited a mean abundance of less than 0.1% **(Fig. 1A & B, Supplementary Fig. 1A & B)**.

**Figure 1.**
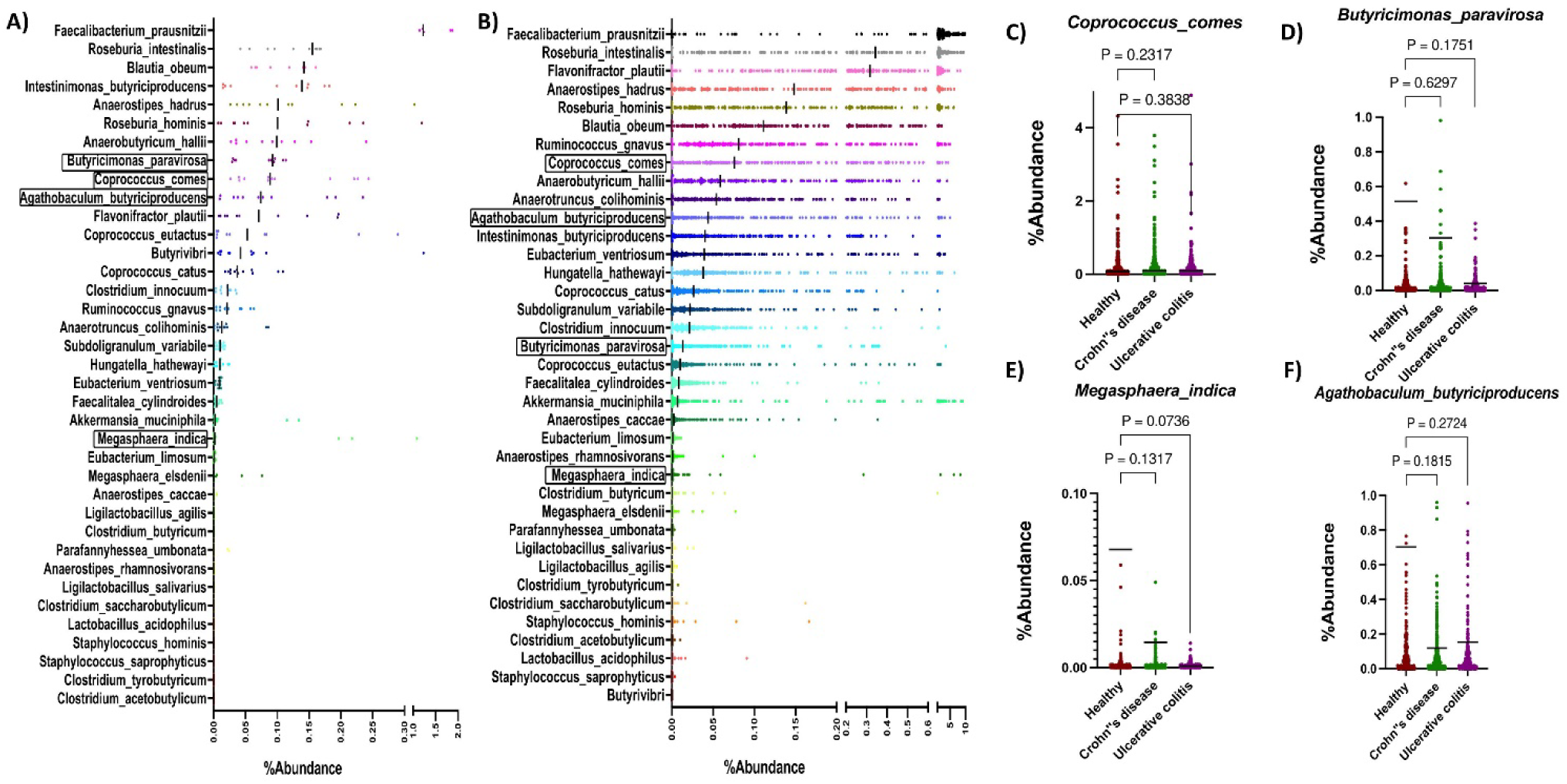
Butyrate-synthesizing bacteria primarily belong to the low dominant category of the human gut. Percentage abundance of Butyrate-producing bacteria in Donor **(A)** and healthy people fecal metagenome samples from a previously published study [61] **(B)**. The highlighted species outlined in box are included in the SC-4 consortia and the dark black line for each species represent the median percentage abundance. The abundance of *Coprococcus comes* **(F)** *Butyricimonas paravirosa* **(G)** *Megasphaera indica* **(H)** *Agathobaculum butyriciproducens* **(I)** in UC, CD patient, and healthy people sample from a previously published study[28]. Comparisons between groups were performed using either the Mann-Whitney t-test. Absolute p-value is represented in the graphs.

The relative abundance of the presumptive butyrate-producing species in our dataset was aligned with the relative distribution across the analyzed public datasets **(Fig. 1B-F)**. The genotypic and phenotypic potential of the selected strains to produce butyrate served as the criteria for their selection. To confirm their genotypic potential, their genome sequences[33] were scrutinized using BLAST to identify markers of the butyrate synthesis pathway, specifically butyryl-CoA: acetate CoA transferase (encoded by *but*) or butyrate kinase (encoded by *buk*). These two genes represent terminal genes in the most prevalent pathway of butyrate production in the gut microbiota[34]. *Butyricimonas paravirosa* and *Coprococcus comes* exhibited the presence of *but*, whereas *Megasphaera indica* and *Agathobaculum butyriciproducens* demonstrated the presence of *buk* in their genome **(Supplementary Fig. 1C)**. Their ability to produce butyrate and other SCFAs was confirmed by *in vitro* culture of the strains in three different media, followed by gas chromatography estimation of SCFAs in the culture supernatant **(Supplementary Fig. 1D)**.

Depletion or enrichment of a bacterial species in a disease condition might indicate the role of that bacterial species in the prevention or exacerbation of the disease, respectively. Nonetheless, this association does not prove their causal relationship with the disease. Thus, to determine the association of selected species with IBD, a disease prevalent in both developed and developing countries [35], the abundance of these bacteria was assessed in fecal metagenomes collected from IBD and non-IBD individuals as part of a longitudinal cohort study conducted by Lloyd-Price et al.[28]. We assessed the abundance of all four species in 598 metagenomes from 50 Chron’s Disease (CD) subjects and 375 metagenomes from 30 Ulcerative Colitis (UC) subjects. We compared it against the abundances of 365 metagenomes from 27 non-IBD subjects. Interestingly, the analysis showed the presence of all four bacterial families under both conditions (healthy and diseased) with a mean abundance of <0.1% **(Fig. 1C-F)**. Compared to the non-IBD microbiota, all four bacteria showed differential abundances in at least one of the IBD conditions. The median abundance of *Agathobaculum butyriciproducens, Megasphaera indica,* and *Butyricimonas paravirosa* was decreased in both UC and CD patients **(Fig. 1D-F)**. We did not find an appealing difference in the median abundance of *Coprococcus comes* in either patients with UC or CD patients. We believe that the lack of statistical confidence is due to the increase in sample size and low abundant nature of the species itself. These results confirmed that there is a reduction in the number of SC-4 bacteria in the gut of IBD Patients, indirectly pointing to the fact that SC-4 bacteria have a negative correlation with IBD condition. Altogether the selected species represented low abundant SCFA producing bacteria shown differential abundance in disease datasets.

### Estimation of intra-consortium interaction of SC-4 *in vivo*

One hypothesis for species being rare within a community is competition with other species. To assess whether the chosen rare species could stably colonize the host gut and to investigate the interactions between these low-abundance species, we employed a germ-free mouse model to polyassociate the SC-4 consortium. After administering equal proportions of the SC-4 mix orally via gavage, fecal samples were collected at different time points to assess the bacterial load. An average bacterial load of >10^7^ CFU/g feces was detected from the 4th day post-inoculation (DPI), which continued to increase until it reached >10^8^ CFU/g feces **(Supplementary Fig. 2A)**. During these time points, we did not observe any significant drop in the bacterial load, suggesting successful and stable colonization of the SC-4 consortium in the mouse gut. An average bacterial count of nearly 10^9^ CFU/g in the cecal content **(Supplementary Fig. 2B)** further verified the successful colonization of these allochthonous host species.

Community profiling via 16s rRNA gene amplicon sequencing revealed the presence of all four bacteria, although their abundance varied and remained consistent throughout the experiment **(Fig. 2A, Supplementary Fig. 2C-E)**. We found a similar community structure in the cecum and colon, but we did not obtain any amplification for 16s PCR from the small intestine contents **(Fig. 2A, Supplementary Fig. 2C)**. This demonstrated the ability of the consortium to colonize the distal gut of germ-free mice. *C. comes* had a higher mean abundance in all mice, whereas *A. butyriciproducens*, *B. paravirosa*, and *M. indica* displayed relatively lower abundance in colonized mice. Next, we examined whether the colonized consortium could produced butyrate and other short-chain fatty acids (SCFAs). As anticipated, the colonized consortium was capable of producing butyrate and other SCFAs. Analysis of cecal content using gas chromatography (GC) detected significantly higher levels of all key SCFAs, except acetate, in SC-4 colonized mice compared than in germ-free mice **(Fig. 2B)**.

**Figure 2.**
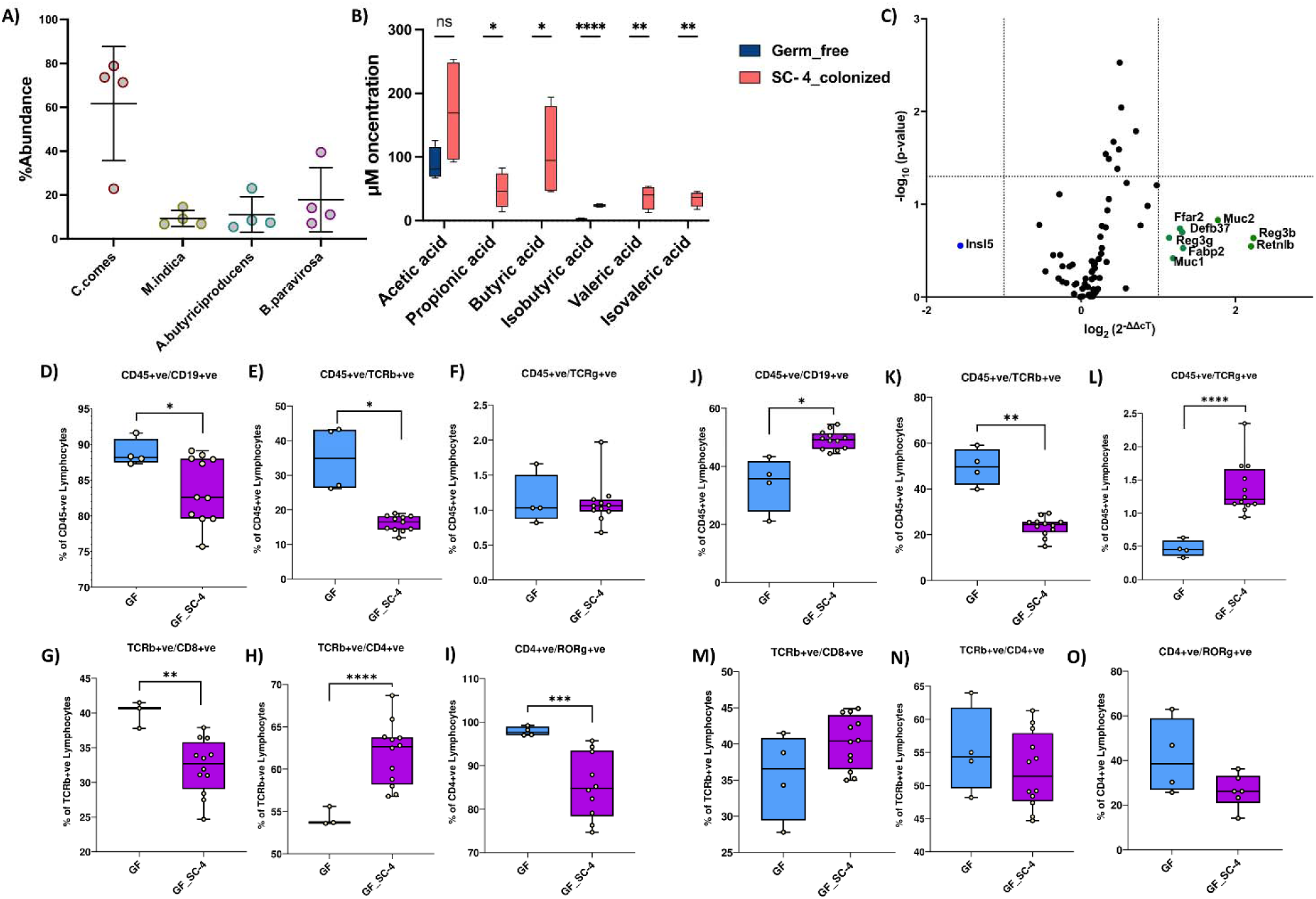
SC-4 successfully colonized germfree mice. Percentage abundance of *Coprococcus comes, Butyricimonas paravirosa, Megasphaera indica, and Agathobaculum butyriciproducens* **(A),** and box plot representing concentration of SCFAs in mice cecum colonized with SC-4**(B)**. RT-PCR gene expression for Rt2 profiler array consisting of 88 genes of mice colon, colonized with SC-4**(C)**. Green dots - overexpressed more than 2-fold change, blue dots-represses more than 2 fold compared to GF control (dotted lines on the x-axis represent a 2-fold increase or decrease in expression and the y-axis represents a p-value cutoff of 0.05). Percentage cell population of lymphocytes in the spleen of GF and GF colonized with SC-4 mice **(D-I)**. Percentage cell population of lymphocytes in the spleen of GF and GF colonized with SC-4 mice **(J-O)**. Comparisons between groups were performed using either the Mann-Whitney U test or the t-test with Welch’s correction. Significance levels were indicated as follows: ns for not-significant, * for p-values < 0.05, ** for p-values < 0.01, and *** for p-values < 0.001.

### Colonization of SC-4 modulates the host immune system

To explore how the consortium interacted with the host, we analyzed the expression profile of a panel of 91 genes in the colon using quantitative Real-Time PCR (qRT-PCR). When compared to germ-free (GF) mice, nine genes showed more than a two-fold change in their expression in the SC-4 colonized group **(Fig. 2C)**. Colonizing GF mice with a potential pathogen usually increases the expression of various pro-inflammatory and anti-inflammatory cytokines, as well as Toll-Like Receptors (TLRs), in the intestine [36] The absence of significant changes in the expression levels of these genes and intestinal barrier proteins in the colon suggests that the SC-4 species do not negatively impact the host, even when colonized at high levels. Notably, changes in expression were found primarily in genes involved in the host’s initial immune defenses, including mucins (MUC-1 and MUC-2) and antimicrobial peptides (Reg3b, Reg3g, Retnlb, and Defb37). Interestingly, SC-4 colonization also increased the expression of fatty acid-binding protein 2 (FABP2/I-FABP) and one of the G-protein coupled receptors (GPRs), FFAR2, or GPR43 **(Fig. 2C)**. FABP2 is a cytosolic protein that binds to Free Fatty Acids (FFA) and is thought to be involved in the uptake and intracellular trafficking of lipids in the intestine. Conversely, FFAR2 is a receptor for short-chain fatty acids [37]. We believe that since the colonized members of SC-4 are capable of producing butyrate in the mouse gut, and butyrate is an energy source for colonocytes, the expression of insulin-like peptide 5 (Insl5) showed more than two-fold downregulation.

To better understand the adaptive immune changes elicited by the colonization of SC-4, we profiled total lymphocytes (CD45+) collected from both secondary lymphoid organs (spleen) and peripheral blood. Flow cytometry analysis of major adaptive immune cell populations, such as B-cells (CD19+), bd-T cells, and subsets of ab-T cells–T-helper (CD4+), T-cytotoxic (CD8+), and Th17 (RORγ+)– showed noticeable differences in the adaptive immune phenotypes between GF and SC-4 colonized mice. Interestingly, the percentage of B cells and ab-T cells in the total lymphocytes was significantly lower in the spleen of SC-4 colonized mice **(Fig. 2D-F)**. Within ab-T cells, there was a significant decrease in CD8+ T cells and increase in CD4+ T cells **(Fig. 2G,H)**. However, the percentage of Th17 cells among CD4+ T cells was significantly reduced in GF mice **(Fig. 2I)**.

In the peripheral blood, ab-T cells showed a similar trend as in the spleen, but there were no significant differences in the proportions of CD4+, CD8+, and Th17 cells compared to GF mice. However, Th17 cells showed a decrease in the proportion of CD4+ T cells, although the difference was not statistically significant **(Fig. 2n)**. In contrast to the spleen, B cells and bd-T cells were significantly increased in BP4 colonized mice **(Fig. 2d-o)**. Histopathological analysis of colon tissue after H&E staining showed normal appearance in both GF- and BPC-colonized gnotobiotic mice without any disruption in mucosal architecture **(Supplementary Fig. 3D)**. However, some morphological changes were observed between the two groups. Both villus length and crypt depth were increased in gnotobiotic mice than in the GF controls. The submucosa and muscularis propria were thicker in gnotobiotic mice than in GF mice, but without any pathological signs such as submucosal edema. We believe this could be related to the increase in expression of MUC-1 and MUC-2 genes when GF mice were colonized with the SC-4 consortium.

### SC-4 colonization protects GF mice from DSS-induced colitis

As the colonization of SC-4 induced positive immune changes in GF mice, we further examined whether this modulation could provide protection against disorders such as IBD. To determine this, SC-4 pre-colonized mice were treated with DSS to induce colitis and monitored throughout the experiment and the mice were sacrificed on day 21 **(Supplementary Fig. 3A)**.The communities formed in the colon and cecum were assessed using 16s rRNA sequencing along with disease status determination using histology assessment of colon samples. Colitis in mice is often characterized by noticeable changes such as shortening of the colon. Interestingly, our results showed that SC-4 pre-colonized mice exhibited less colon shrinkage than the control group **(Fig. 3A&B)**, suggesting that SC-4 pre-colonization may offer systemic protection against DSS-induced colitis. SC-4 colonization also protected mice against colitis-induced death **(Supplementary Fig. 3B)**. Histopathological analysis of GF mice with colitis showed moderate inflammation, loss of goblet cells, vascular proliferation, and wall thickening ; however, mice pretreated with SC-4 showed no to mild inflammation in most samples as indicated by the decrease in pathology score **(Fig. 3C, Supplementary Fig. 3C)**. Since we did identify substantial differences in MUC-1 and MUC-2 gene upon SC-4 colonization, we hypothesized that the protective effect conferred by SC-4 might be attributed to the improvement of gut barrier function. These genes, which contribute to the thickness of the submucosa and muscularis propria, may enhance barrier function. PAS staining on the histology sections showed a significant increase in the number of goblet cells present confirming the hypothesis **(Fig. 3C&E)**. The 16s rRNA sequencing of the feces showed the same trend as that previously observed for SC-4 colonization **(Fig. 3F)**. *C. comes* relative abundance slightly decreased with an increase in abundance for the rest of the three species **(Fig. 3F)**. This suggests that *C. comes* may have an antagonistic effect on the other three species. This trend remained true for both the samples taken from colon and the cecum **(Fig. 3F, Supplementary Fig. 3C)**. Furthermore, quantification of short-chain fatty acids showed no difference when treated with DSS **(Fig. 3G)**. A slight increase in butyrate production was also observed. This implies that the induction of a disease state does not reduce SC-4 ability to produce SCFAs.

**Figure 3.**
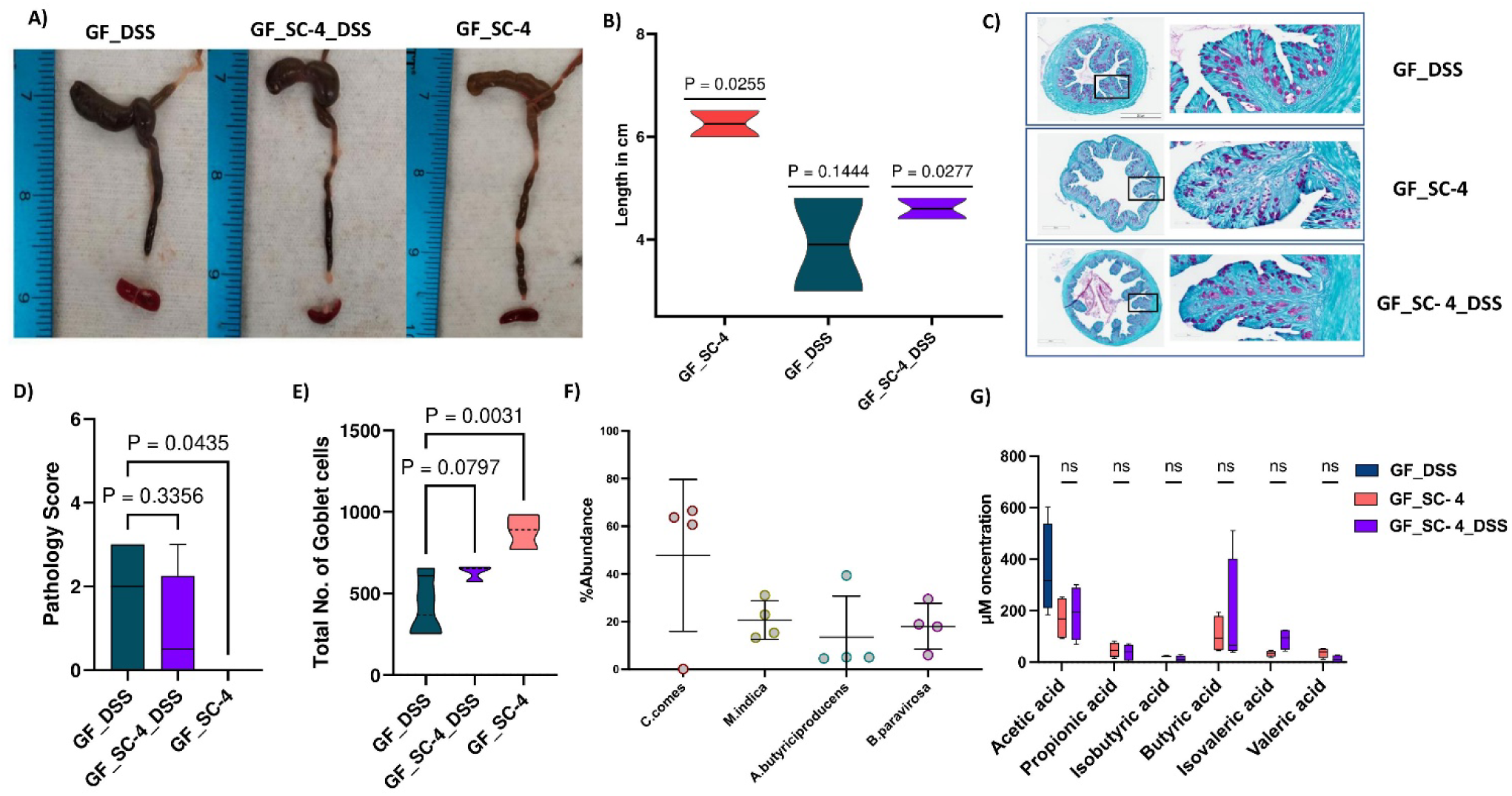
SC-4 confers protection against DSS-induced colitis. Representative colon from Germ free (GF) mice induced with colitis (GF_DSS), GF mice gavaged with SC-4 bacteria (SF_SC-4) and GF mice gavaged with SC-4 bacteria induced with colitis (GF_SC-4_DSS) **(A)** and associated Violin plot **(B)** representing the length of the colon in cm. Statistical significance is calculated using one sample t-test across across the complete population and the absolute value for each group is represented above each violin plot. Representative histopathological photograph showing the colon tissue cross-section after PAS staining GF_DSS, GF_SC-4 and GF_SC-4_DSS mice groups **(C).** Box plot mentioning the pathology score calculated from the histopathological section of colon tissue cross section after H&E staining as mentioned in (Supplementary Fig.3D) **(D)**. Truncated Violin plot representing total number of goblet cells counted from the histopathological section of colon tissue cross section after PAS staining as mentioned in (c) **(E)**. Statistical significance for D & E calculated using one-way annova with Dunnet correction with absolute p-value represented on the graph. Percentage abundance of SC-4 bacteria **(F)** and box plot representing concentration of SCFAs **(G)** in mice cecum upon DSS treatment. Comparisons between groups were performed using either the Mann-Whitney U test or the t-test with Welch’s correction.

Furthermore, we examined the differential gene expression upon disease induction for selected genes from the previous panel **(Fig. 2C, Supplementary Table. 1)**. However, compared with the control, SC-4 mice treated with DSS did not show any biologically relevant changes **(Fig. 4A & B)**. We also checked for the systemic immune response; apart from ab-T cells, there were no significant changes in the cell profile in DSS-treated animals **(Fig. 4C-N)**. There was a substantial reduction in the ab-T cell population in both PBMC and spleen of SC-4 pre-colonized mice **(Fig. 4D, J)**. This result is comparable to that previously observed when GF mice were colonized with SC-4 **(Fig. 3J, M)**.

**Figure 4.**
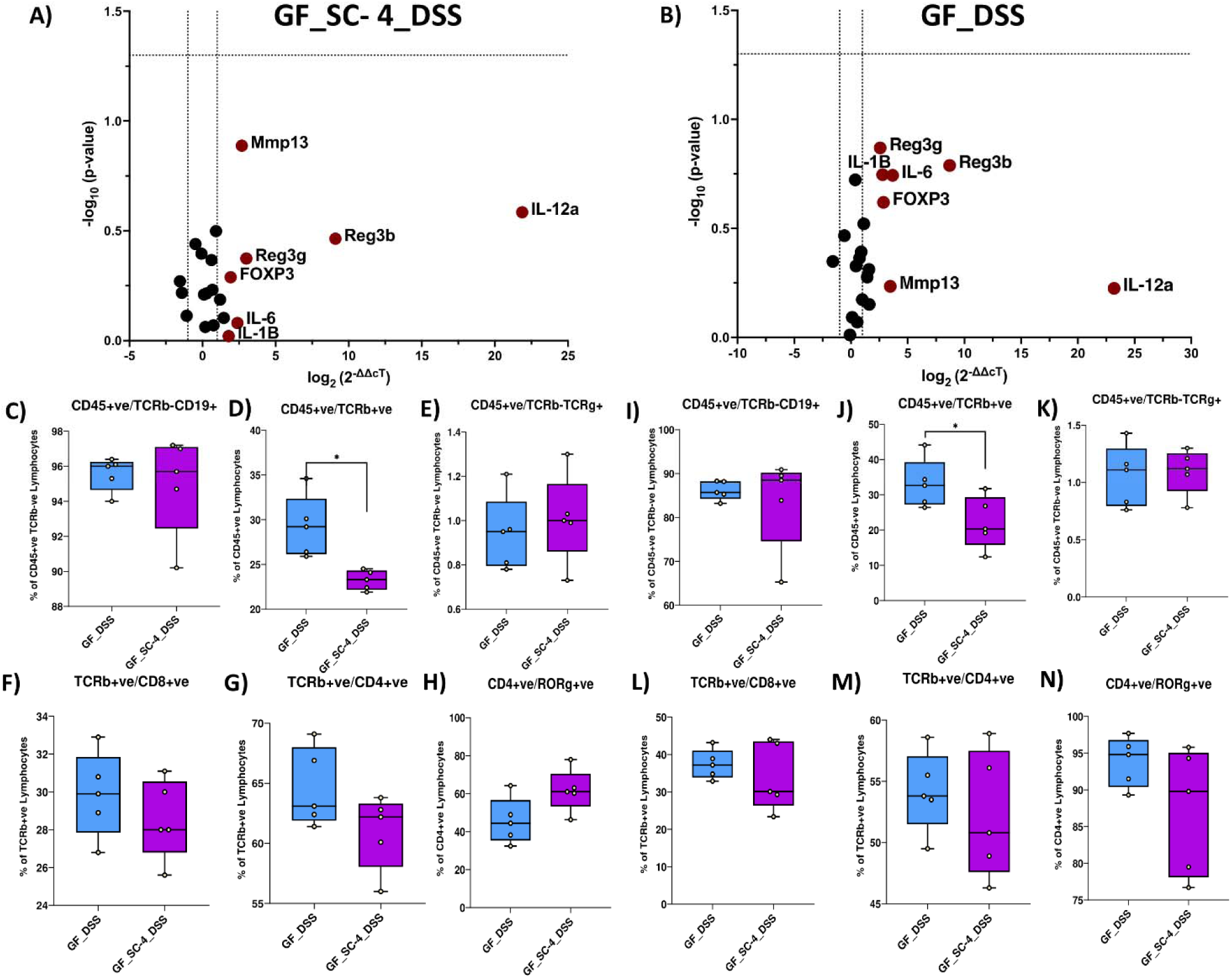
Local and systemic immune response in colitis model of GF mice treated with SC-4. RT-PCR gene expression for selected 25 immune-related genes of mice colon, GF mice pre colonized with SC-4 and then induced with colitis (GF_SC-4_DSS) **(A)** and GF mice induced with colitis (GF_DSS) compared to GF control **(B)**. Green dots - overexpressed more than 2-fold change, Red dots - overexpresses more than 2 fold compared to GF control (dotted lines on the x-axis represent a 2-fold increase or decrease in expression and the y-axis represents a p-value cutoff of 0.05). The percentage cell population of lymphocytes in the spleen of GF and GF colonized with SC-4 induced with colitis **(C-H)**. The percentage cell population of lymphocytes in the spleen of GF and GF colonized with SC-4 mice induced with colitis **(I-N)**. Comparisons between groups were performed using either the Mann-Whitney U test or the t-test with Welch’s correction. Significance levels were indicated as follows: * for p-values < 0.05, ** for p-values < 0.01, and *** for p-values < 0.001.

However, no significant TH17 cells was increased in the spleen nor PBMC of GF mice with colitis. Our assumption for this change could be attributed to the previous recognition of the molecular patterns of the SC-4 consortium. Taken together, SC-4 consortium protect against DSS induced colitis in GF mice by improving the mucus barrier function.

### SC-4 members show a negative correlation with colitis in Humanized mice

Next, we examined whether the SC-4 consortium could colonize mice, establish a higher abundance, modulate microbiome composition, and improve host health in the presence of complex human microbiota. To this end, we gavaged the SC-4 consortium in human fecal microbiota-colonized mice. The composition of the microbiome following SC-4 gavage was analyzed by 16s rRNA sequencing of the fecal pellets. While *Butyricimonas paravirosa* (Odoribacteraceae)*, Coprococcus comes* (Lachnospiraceae) and *Agathobaculum butyriciproducens* (Ruminococcaceae) were detected in the community **(Fig. 5A, 5H)**, *Megasphaera indica* (Veillonellaceae) was not detected in any of the mice. This indicates that these three bacteria can colonize the mouse gut, even when introduced as part of a complex community. Furthermore, we examined whether the abundance of these three species changes during colitis. To this end, we induced colitis in SC-4 colonized humanized mice.

**Figure 5.**
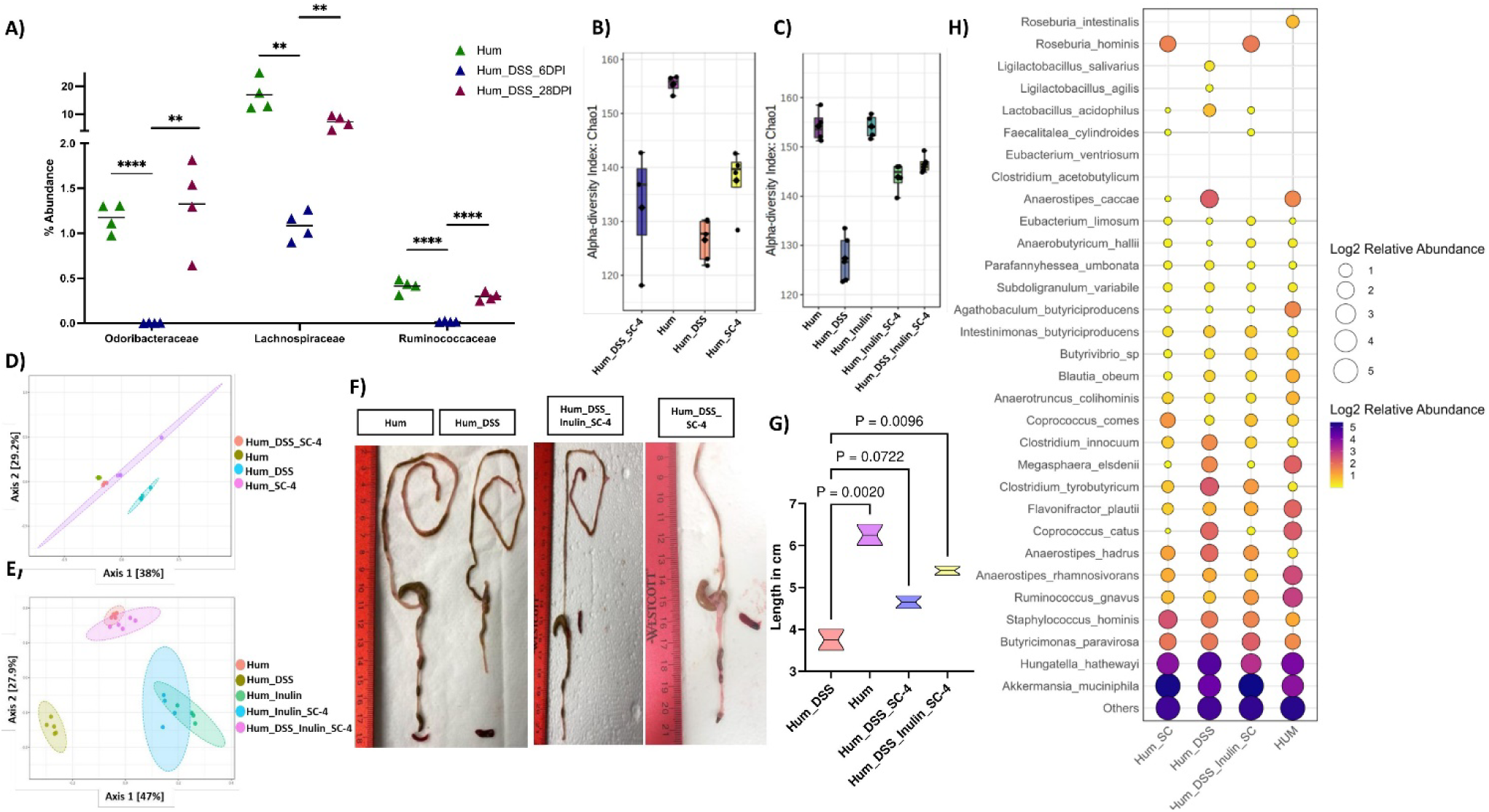
SC-4 colonization increases Microbiome diversity. Percentage abundance of *Butyricimonas paravirosa* (Odoribacteraceae), *Coprococcus comes* (Lachnospiraceae), or *Agathobaculum butyriciproducens* (Ruminococcaceae) in humanized mice (Hum), humanized mice induced with colitis (Hum_DSS) at 6^th^ and 28^th^ Day post-infection **(A)**. Box plot representing alpha diversity analysis **(B, C)** and PCoA representing bneta diversity analysis **(D, E)** (Chao1) when humanized mice with DSS-induce colitis treated with SC-4 (Hum_DSS_SC-4), Humanized mice gavage with SC_4 (Hum_SC-4), Hum_DSS, Hum **(B,D)**, humanized mice treated with SC-4 with supplementation of Inulin (Hum_Inulin_SC-4), humanized mice with DSS-induce colitis treated with SC-4 with supplementation of Inulin (Hum_DSS_Inulin_SC-4) and humanized mice with supplementation of Inulin (Hum_Inulin) **(C,E)**. Representative colon images from Hum, Hum_DSS, Hum_DSS_SC-4, and Hum_DSS_Inulin_SC_4 **(F)** and associated Violin plot **(G)** representing the length of the colon in cm. Statistical significance for (G) calculated using one-way annova with Dunnet correction with absolute p-value represented on the graph. Bubble plot representing relative abundance in natural log scale at a species level resolution for SCFA producing bacteria (Similar to Figure 1B) from Hum, Hum_DSS, Hum_DSS_SC-4, and Hum_DSS_Inulin_SC_4 mice groups **(H)**.

The abundance of all three species was significantly reduced on the 6th day post-induction **(Fig. 5A)**. However, the reduction in abundance reverted significantly on the 28^th^ day post-gavage when the mice started to recover from colitis **(Fig. 5A)**. This indicated a negative correlation between these three species and colitis in mice. As the abundance of these bacteria increased with time post-gavage, we hypothesized that the introduction of these species back into the system could help accelerate recovery from DSS-induced colitis **(Supplementary Fig. 4A)**.

We examined the microbiome composition following these experiments and found that the introduction of SC-4 increased the alpha diversity of the gut community in DSS-treated mice, whereas it was lower in control mice **(Fig. 5B, Supplementary Fig. 5A-D)**. When the beta diversity between the different groups was examined, we observed that SC-4 administration after colitis induction shifted the diversity to resemble that of healthy humanized mice **(Fig. 5B, Supplementary Fig. 5E-H)**. This suggests that introducing SC-4 can increase diversity and shift the microbiome to a controlled state. Besides the fact that SC-4 increased diversity, it also increased colon length, suggesting a higher rate of recovery for DSS-induced colitis **(Fig. 5F&G)**.

As fermentation of dietary fiber has been shown to promote the abundance of SCFA-producing species, we also examined whether the addition of dietary fiber could increase the abundance of SC-4 and provide better protection against colitis. To this end, the above set of experiments was repeated with and without inulin, a commonly used dietary fiber. Our analysis found that the addition of inulin increased the alpha diversity at a much higher rate **(Fig. 5C)**. When beta diversity was analyzed, we found that colonization of SC-4 with Inulin treated group clustered more towards control mice **(Fig. 5E)**. However, inulin and inulin with SC-4 administration in control mice had a different clustering, which was expected because of the effect of inulin on the microbiome **(Fig. 5C)**. This also proves that inulin alone is less efficient in increasing diversity compared to SC-4 supplementation. Further addition of inulin with SC-4 resulted in an increase in colon length compared to treatment with SC-4 alone **(Fig. 5F &G)**. Moreover both SC-4 and Sc-4 with Inulin treatment showed a decrease in diarrhea in mice as represented as fecal score 8 and 9 post-induction respectively **(Supplementary Fig. 5I)**. Because we know that a stable microbial community is functionally redundant, we wanted to check whether DSS treatment showed this redundancy in humanized mice. To this end, we created a custom database of full length 16s rDNA for the known SCFA-producing bacterial species, as shown in **(Fig. 1A-B)** and checked their abundance in various treatments. This analysis showed a decrease in most of the highly abundant SCFA-producing bacteria upon DSS administration, while low-abundant bacterial species such as *Clostridium innocuum, Anaerostipes hadrus, Clostridium tyrobutyricum, and Hungetella hathewayi* increased in abundance **(Fig. 5H)**. Furthermore, in the case of SC-4 and SC-4 along with inulin treatment, the supplementation appeared to increase in the rest of the SCFA producers in the community **(Fig. 5H)**. Interestingly, in both treatments, we also found an increase in the abundance of *Akkermansia mucicniphila* a known butyrate-producing species with known gut health-improving functions. This suggests that SC-4 administration might help regain SCFA production through interspecies interactions that boost other beneficial species.

We checked gene expression changes in the mouse colon upon colonization of humanized mice with the SC-4 consortium post euthanasia (day-23). As noticed in the case of GF mice, we did not observe any significant changes in inflammatory-related gene expression, further proving the non-pathogenic nature of the SC-4 consortium **(Supplementary Fig. 5J-L)**. An iNOS-dependent increase in colonic mucus has previously been demonstrated in colitis-induced rats. We observed a significant increase in iNOS expression when colitis-induced mice were treated with SC-4 or SC-4 supplemented with inulin **(Fig. 6A-C)**. We believe that SC-4 might be helping the mice to protect against more damage from colitis by increasing mucus production through the increase of iNOS. There was also a reduction in the expression of IL-6 compared to colitis recovered naturally. Myeloperoxidase (MPO) produces hypohalous acid to carry out its antimicrobial activity and is upregulated in colitis recovered naturally **(Fig. 6A)**. This might be due to the perturbation and community shift due to DSS-induced colitis, which increased the abundance of some pathogenic bacterial classes and, hence, the response. The same was downregulated when the mice were treated with SC-4 or SC-4 supplemented with inulin. This indirectly shows the ability of the consortium to transform the community into a healthy state. Histopathology of the colon sections showed several differences between BP-treated and untreated mice. Colitis-induced humanized mice **(Supplementary Fig. 6A)** showed a phenotype similar to that of GF mice with colitis, whereas when treated with SC-4 or SC-4 supplemented with inulin **(Fig. 6H, Supplementary Fig. 6B)** showed a phenotype similar to that of control mice. The Pathology score was more than 0 for only humanized mice induced with colitis and not for any of the treatment **(Fig. 6H)**. PAS staining of the histology section confirmed the our observation that iNOS associated goblet cell increase. We saw an increase in the goblet cell number for both SC-4 and SC-4 and inulin treatment, but significant for Inulin and Sc-4 treatment only **(Fig. 6D-G & I)**. These results suggest that SC-4 with dietary fibers supplementation may help restore the dysbiotic microbiome.

**Figure 6.**
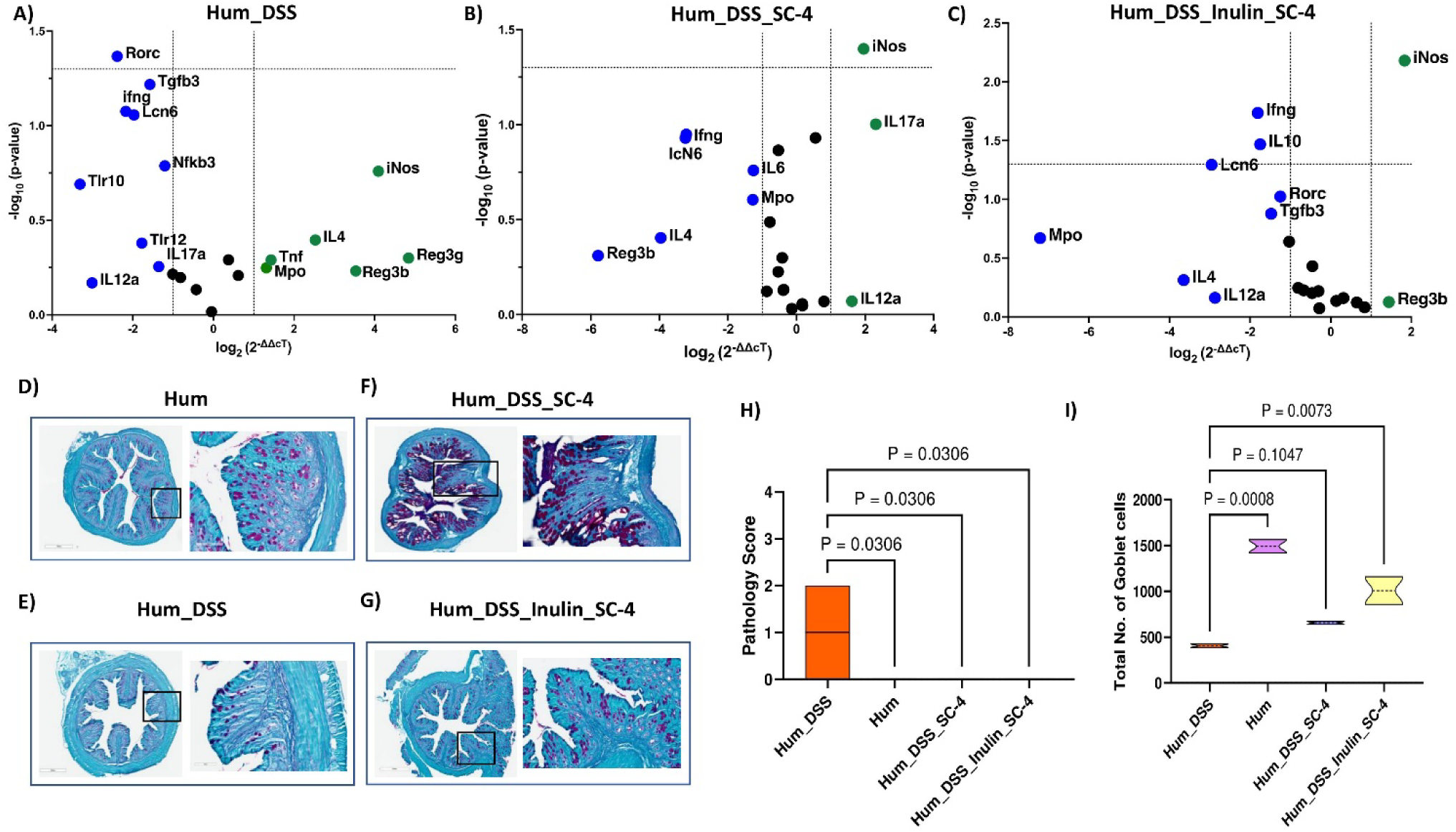
SC-4 colonization in Humanized mice increases iNOS gene expression. RT-PCR gene expression for selected 25 immune-related genes of mice colon represented in volcano plot for humanized mice with DSS-induce colitis (Hum_DSS) **(A),** humanized mice induced with colitis and treated with SC-4 (Hum_DSS_SC-4) **(B),** and humanized mice induced with colitis and treated with SC-4 and supplemented with Inulin (Hum_DSS_Inulin_SC-4) compared to humanized mice control (Hum)**(C)**. Green dots - overexpressed more than 2-fold change, green dots- overexpresses more than 2-fold compared to HUM control (dotted lines on the x-axis represent a 2-fold increase or decrease in expression and the y-axis represents a p-value cutoff of 0.05). Representative histopathological photograph showing the colon tissue cross-section after H&E staining for Hum**(D),** Hum_DSS **(E),** Hum_DSS_SC-4 **(F),** Hum_DSS_Inulin_SC-4 **(G)** mice groups. Pathology score calculated from the histopathological section of colon tissue cross section after H&E staining as mentioned in (Supplementary 6A,B) **(H)**. Total number of goblet cells counted from the histopathological section of colon tissue cross section after PAS staining as mentioned in (D-G) **(I)**. Statistical significance for H & I calculated using one-way annova with Dunnet correction with absolute p-value represented on the graph.

## Discussion

In various biological systems such as terrestrial, aquatic, and animal gut microbiomes, a skewed distribution of species abundance is often observed. Within the healthy human gut microbiome, this phenomenon is particularly pronounced; while hundreds of species exist, only a few are present in high abundance, and the vast majority are found in low abundance. Researchers refer to these low-abundant species as the “rare biosphere” of microbiomes[38]. This rarity can be visualized in a rank abundance plot, where the long tail represents these rare species after arranging them from high to low abundance. A challenge in studying this area is the lack of a universal cutoff for defining rarity. Most studies arbitrarily considered species with less than 0.1% abundance as rare or low abundant. Applying this threshold revealed that over 95% of the species in the human gut microbiome fit this category. Despite this prevalence, most microbiome research has concentrated on the functional role of high-abundance species, leaving the role of rare species relatively unexplored.

In this study, we investigated the role of four low-abundant gut bacterial species, *B. paravirosa, C. comes, M. indica,* and *A. butyriciproducens*, on host health. These were chosen for their ability to produce short-chain fatty acids (SCFAs), a trait that we focused on because of SCFAs’ importance in gut homeostasis. More specifically, these selected species were found to be low-abundant and possessed genes responsible for producing butyrate, a major SCFA with demonstrated positive effects on gut health. In our search for the abundance of these bacteria within a dataset of 598 metagenomes, which included samples from 50 subjects with Crohn’s Disease (CD), 30 subjects with Ulcerative Colitis (UC) comprising 375 metagenomes, and 27 non-IBD subjects with a total of 365 metagenomes, we observed a decrease in the median abundance of all the bacteria in cases of UC or CD. However, this reduction was not statistically significant. Several studies have revealed that butyrate-producing genera such as *Alistipes, Barnesiella, Faecalibacterium, Oscillibacter, Agathobacter,* and *Ruminococcus* are depleted in patients with Inflammatory Bowel Disease (IBD) [39]. Additionally, results from randomized controlled trials have shown that Fecal Microbiota Transplantation (FMT) can replenish lost diversity and ameliorate UC symptoms in affected patients [40, 41]. However, the efficacy of defined bacteriotherapy or probiotic therapy in treating CD remains inconclusive [42, 43]. This uncertainty might be attributed to the fact that most interventions have been carried out using high-abundant taxa such as Bifidobacteria and Lactobacilli [44]. The analysis was conducted using both germ-free and human microbiota-associated mouse models. We formed a synthetic consortium (SC-4) with the selected species and closely monitored their impact on host health, laying the groundwork for potential therapeutic applications targeting low-abundant, SCFA-producing gut bacteria.

Several traits common to high-abundance SCFA-producing species were also observed in the species examined in our study. Specifically, SC-4 colonized germ-free mice in large numbers and produced SCFAs **(Fig. 2)**. Importantly, the colonized mice displayed no signs of infection or pathology, indicating the safety of these strains. The primary effect of SC-4 colonization in germ-free mice was an increase in the expression of the MUC-1 and Muc2 genes and an increase in mucin thickness in the colon. These results resemble those found in the treatment with sodium butyrate, which quadrupled ex vivo mucin synthesis in colonic biopsy specimens [45]. Similarly, treating human-polarized goblet cell lines with butyrate as an energy source augmented the expression of various MUC genes [46]. SC-4 colonization also conferred protection against DSS-induced colitis **(Fig. 3)**.

Given that the protective efficacy observed in a germ-free animal model might not translate to a more complex microbiota environment, we tested SC-4’s effectiveness using a human microbiota-associated mouse colitis model. Surprisingly, SC-4 not only cured the mice with colitis but also increased the microbiome diversity to a healthier state **(Fig. 5-6)**. *M. indica,* however failed to colonize in this intricate setting. We noticed a similar increase in mucin thickness but did not observe any differences in MUC gene expression. Instead, a notable increase in iNos gene expression was detected, reminiscent of the iNos-mediated growth in colonic mucus observed in a colitis rat model [47]. We hypothesized that the same mechanism might apply to humanized mice for protection and cure. While the observed increase in iNOS expression associated with SC-4 treatment suggests a possible mechanism for enhanced mucus production, this hypothesis necessitates additional mechanistic studies for validation. There were no significant changes in the adaptive immune response in either germ-free or humanized mice.

The inability of SC-4 to show any visible differences in the adaptive immune profile and major signaling pathways compared to DSS-treated mice prompted us to ask whether the observed phenotype was associated with functional redundancy maintained by the humanized mouse gut community. We also found that the functionality of SCFA production was retained by the community, with an increase in some of the low-abundant SCFA producers after DSS treatment **(Fig. 5H)**. This is experimental proof for the insurance hypothesis being in existence for gut microbiome communities, similar to what has been reported for other biotic communities. This hypothesis extends from the broader theory that biodiversity sustains ecosystem stability by enabling alternative species to take over functional roles if others fail under altered conditions. Applied to the microbiome, this diversity enables the microbial community to adapt to changes such as diet alterations, infections, or antibiotic treatments, continuing to provide vital functions, such as nutrient assimilation, immune system regulation, and resistance to pathogens[48]. This adaptive quality serves as an “insurance” against disruptions that could lead to dysfunction or illness. This phenomenon vanished when the system was introduced with SC-4. This poses important questions in microbiome research: Do prevalence studies identifying specific bacterial species presence in different physiological states correspond to this insurance species? Can we categorize a bacterial species to be associated with a disease by a mere increase in abundance in the gut community during the disease course? These questions must be answered before categorizing any bacteria that are positively associated with a disease. For instance, without considering this hypothesis, one can argue that the increased abundance of species in this study has a positive association with IBD.

Altogether, our findings enhance the fundamental understanding of the functions of low-abundant species and provide a proof-of-principle study for exploring these species as potential probiotic candidates. These results align with the biological insurance hypothesis within the gut microbial ecology [49]. Specifically, in the context of Inflammatory Bowel Disease (IBD), low-abundant species we tested may assume the roles of depleted high-abundant taxa such as *Alistipes, Barnesiella, Faecalibacterium, Oscillibacter, Agathobacter, and Ruminococcus* [39]. Further population-level studies are required to confirm this possibility. The future research on the SC-4 bacterial consortium as a treatment for IBD will focus on understanding its ecological role on administration across diverse fecal microbiomes and how diet influences its therapeutic effectiveness. Key efforts will involve investigating the differential impacts of SC-4 and other low abundant SCFA producing microbiome members on metabolomic profiles in varied microbiome backgrounds, aiming to tailor treatments to individual gut ecologies. This approach promises to enhance the precision of microbiome-based therapies by incorporating patient-specific microbial compositions and dietary interactions, thereby optimizing the potential benefits of SC-4 in managing IBD. Our present study paves the way for the exploration of low-abundant members in maintaining gut diversity and developing novel therapeutics against colitis.

## Materials and Methods

### Animal experiments

Six-week-old C57BL/6 Germ-free (GF) mice were procured from Taconic Biosciences Inc. ( New York, USA) and the Mayo Clinic (Rochester, USA). To negate any bias associated with gender differences, we ensured the inclusion of both male and female mice in our experiments, where feasible. All animals were subjected to a 12-hour light/dark cycle and provided with unlimited access to sterile drinking water and a standard chow diet. The protocols for all animal experiments were reviewed and approved by the South Dakota State University (SDSU) Institutional Animal Care and Use Committee (approval #19-014A).

### Bacteria culture and maintenance

The bacterial species utilized in our experiments were cultured and maintained using DSMZ modified PYG media, supplemented with L-cysteine as a reducing agent and resazurin as an oxygen indicator. The cultures were preserved at 37°C in an anaerobic chamber (Coy Lab Products Inc., MI, USA) filled with an atmospheric composition of 85% nitrogen, 10% carbon dioxide, and 5% hydrogen.

For *in-vitro* phenotypic assessment of butyrate production, overnight bacterial cultures were prepared in three distinct media conditions. These included Brain Heart Infusion broth and yeast extract-based media, with and without inulin as a complex carbohydrate supplement. The third medium used was DSMZ modified PYG medium. After 24 h of incubation in the anaerobic chamber, 100µL of the culture was combined with 500µL of 5% freshly prepared meta-phosphoric acid. This mixture was then thoroughly vortexed for two minutes and promptly stored at -80°C for subsequent gas chromatography (GC) analysis, after which the optical density (OD_600_) of overnight grown cultures was adjusted to one, and aliquots were stored in 12% DMSO at -80°C until further use. On the day of inoculation, an equal volume of all four bacteria were resuspended and washed twice in the PYG medium. After centrifugation at 7000 RPM for 10 min, the bacterial pellets were reconstituted in PYG medium to get a 20x concentrated bacterial suspension. 100uL / mice of bacterial suspension were prepared and transferred to GF mice units in sterile, airtight 2mL glass vials.

### Mice experiments

In our experiments involving gnotobiotic mice, we utilized a mixture of four butyrate-producing bacterial strains, referred to as the ‘Short-chain fatty acid producing consortium” (SC-4), for inoculation. A 200µL dose of this 20X concentrated bacterial mixture was administered to the treatment group via oral gavage. Each experimental group consisted of to 4-6 mice, with an additional group of uninoculated mice serving as the germ-free control. Each mouse received two inoculations (once per day) and was subsequently allowed a colonization period ranging from a minimum of 14 days to a maximum of 28 days. At the conclusion of the colonization period, all mice were euthanized via cervical dislocation and samples were collected for subsequent analysis.

Colitis was chemically induced in germ-free (GF) mice following the protocol described by Wirtz et al. [50], with slight modifications. Dextran sulfate sodium salt (30-50 KDa) from MP Biomedicals, USA was added at a concentration of 1.5% to autoclaved drinking water. Mice had ad libitum access to DSS-infused water for six days, after which they were euthanized via cervical dislocation, and samples were subsequently collected for further analysis. On the fifth day, all animals displayed signs of rectal bleeding. One mouse in the GF group treated with DSS died on the sixth day of treatment. The onset of colitis was validated using a fecal occult test conducted on the third day.

In the case of human Fecal Microbiota Transplant (Hum) mice, both male and female six-week-old mice received oral gavage of 200µL of the pooled fecal sample over three consecutive days, as described in our previous work [33]. After several generations of breeding, male and female mice were exposed to 2.5% Dextran Sulfate Sodium (DSS) in their drinking water for five days to induce colitis. The mice were then switched to regular drinking water for another three days before being sacrificed in the Hum_DSS group post 23days. For the Hum_DSS_SC-4 and Hum_DSS_SC-4_Inulin groups, mice were gavaged with 200µL of a 20X concentrated bacterial mixture for two days, or the bacterial mix along with 1% inulin in their drinking water. In the Hum_SC-4, Hum_SC-4_Inulin, and Hum_Inulin groups, humanized mice were treated with the bacterial mix, bacterial mix along with inulin, or inulin only, and were then sacrificed after 23 days. Cecal and colon fecal samples were collected for 16S rRNA amplicon sequencing.

### Fecal occult blood test

The initiation of colitis was confirmed by an occult blood test using fecal samples. Whatman filter paper pre-treated with guaiac was used for this test. A thin smear of feces was applied to one side of a guaiac-coated Whatman filter paper. Subsequently, one or two drops of 3% hydrogen peroxide were carefully administered on the opposite side. A rapid transition to blue on Whatman paper was interpreted as a positive indication of the presence of blood in the sample.

### Bacterial enumeration

The total bacterial count in fecal samples collected at different time points was assessed to evaluate bacterial colonization. Anaerobically modified PYG medium (DSMZ medium) was used to enumerate the bacterial load, as this medium was found to support all four bacteria used in this study.

### Genomic DNA extraction and targeted amplicon sequencing

Genomic DNA was extracted from pure bacterial cultures using a DNeasy blood and tissue kit (Qiagen, Maryland, USA). We used the DNeasy power soil kit (Qiagen, Maryland, USA) to extract fecal samples and intestinal contents of the mice, according to the manufacturer’s instructions. The 16s rRNA gene sequencing was performed according to the standard Illumina protocol, where PCR amplicons targeting the V3-V4 region of the bacterial 16s rRNA gene were used for sequencing. Locus-specific primer pairs (16S Amplicon PCR Forward Primer = 5’ CCTACGGGNGGCWGCAG, and 16S Amplicon PCR Reverse Primer = 5’ GACTACHVGGGTATCTAATCC) were attached to overhang adaptors (Forward overhang:5’ TCGTCGGCAGCGTCAGATGTGTATAAGAGACAG, and Reverse overhang:5’ GTCTCGTGGGCTCGGAGATGTGTATAAGAGACAG) at the 5’ end of the respective primer sequences (Illumina, Inc.) and used to amplify the region of interest.

The resulting amplicons were then cleaned of free primers and primer dimer species using AMPure XP beads (Beckman Coulter). Sequencing libraries were prepared using the Nextera XT library preparation kit (Illumina, Inc.). After indexing using a dual barcoding system according to the manufacturer’s protocol and then normalizing to a concentration of 4nM, the libraries were pooled into a single loading library. Sequencing was performed on an Illumina MiSeq platform using 2 × 300 bp paired-end read chemistry. Demultiplexed and adaptor-trimmed reads generated by Illumina analytical tools were used for further processing.

### Bacterial community profiling

Microbiota profiling from 16S sequencing was performed using Vsearch [51]. Merging and Quality filtering with a minimum cutoff length of 400 and maximum length of 500 were performed for the fastq files using the Vsearch tool. Singleton and chimeric reads (UCHIME) were removed. OTU selection was performed using VSEARCH abundance-based greedy clustering. OTUs were annotated using the SILVA reference database [52] or Custom database made from full-length 16s extracted from the whole genome of the four bacteria in the consortia. To create a custom database for the SCFA-producing bacterial consortium, we used the full length 16s rRNA sequence obtained from SILVA/NCBI. The resulting OTU table was further processed for analysis. Alpha and Beta diversity calculations and plotting were performed using Microbiome Analyst with the default parameters [53].

### Bacterial species abundance mapping

To assess the association of the four selected butyrate producers in IBD conditions, an abundance mapping at the strain level was performed using publicly available 1338 gut metagenome sequencing datasets collected from US individuals as a part of a longitudinal cohort research study by Lloyd et al.[54]. For all metagenome reads, quality trimming and adapter clipping were performed using Trimmomatic [55]. Furthermore, the reads were aligned against the human genome to filter out human reads and assembled using Bowtie2 v2.3.2 [56] and SAMtools [57]. The resulting contigs were classified taxonomically by k-mer analysis using Kraken2 [58] with the Kraken 2 standard bacterial database built using Kraken-built. The subsequent estimation of species abundance was performed using the Kraken tools [59].

### Gas chromatography (GC) analysis for SCFAs

Immediately before the GC analysis, the samples were thawed and vortexed for 2 min. The ethyl acetate (EA) extraction method described by Garcia-Villalba et al. [60] was used to detect SCFA. Briefly, the homogenized sample was centrifuged for 10 min at 17,949 × g. Each milliliter of supernatant was extracted with 1 mL of an organic solvent (EA) for 2 min and centrifuged for 10 min at 17,949 × g. A minimum of 200uL organic phase volume per sample was loaded in the Trace 1300 gas chromatogram (Thermo Fisher Scientific, USA).

### Quantitative RT-PCR-based Gene expression profiling

To assess the localized response of host-microbe interactions following colonization of BP4 strains, 100 mg of tissue from the colon was sampled and snap-frozen in liquid nitrogen. The samples were then stored at -80 °C until further processing. For RNA extraction, TRIzol® (Ambion, Life Technologies, USA)-chloroform (Sigma-Aldrich, USA) was used. DNase treatment was performed using an RNase-Free DNase kit (Qiagen, Maryland, USA) according to the manufacturer’s protocol to remove any contaminant genomic DNA. RNA quality and quantity were evaluated using a NanoDrop One (Thermo Scientific, USA) and stored at -80°C until further use.

Complimentary DNA (cDNA) was prepared from 250 ng of total RNA using the First Strand cDNA Synthesis Kit (New England BioLabs Inc., USA), according to the manufacturer’s protocol. Host gene expression was assessed by qRT-PCR using an Rt2 profiler array (Qiagen, Maryland, USA) **(Supplementary. Table1)** or 25 defined immune-related genes **(Supplementary. Table2)**. Reactions were prepared using the Power SYBR® Green PCR Master Mix (Applied Biosystems, USA). PCR was performed in an ABI7500 standard (Applied Biosystems, USA) RT-PCR machine under the following cycling conditions: 95 °C for 10 min, 40 cycles of 95 °C for 15 s, and 60 °C for 1 min.

Raw cycle threshold (C_T_) values at a threshold of 0.15 were exported and then uploaded to the Qiagen data analysis center for further analysis. A C_T_ cutoff value of 30 was used for the analysis. The average geometric mean of the C_T_ values of two housekeeping genes, mouse beta-actin (Actb) and mouse glyceraldehyde phosphate dehydrogenase (Gapdh), were used for data normalization and ΔC_T_ calculation. The fold change and fold regulation of gene expression were calculated using the ΔΔC_T_ method.

### Separation of lymphocytes from Peripheral mouse blood

Immediately after euthanasia, blood was collected via cardiac puncture, transferred to heparinized blood collection vials (BD Vacutainer), and gently mixed to prevent coagulation. SepMate TM PBMC isolation tubes (STEMCELL Technologies, Canada) were used to collect lymphocytes from blood. An equal volume of DPBS (Dulbecco’s PBS) was added to the blood sample before transferring to SepMate TM tubes prefilled with 4.5 mL of Lymphoprep (STEMCELL Technologies, Canada) density gradient medium. The tubes were then centrifuged at 1200 × g for 10 min at room temperature. The top layer containing the plasma and mononuclear cells (MNC) was transferred to a fresh 15mL falcon tube by a single pour-off step. After washing twice with DPBS, the transferred MNCs were re-suspended in freezing media containing 10% DMSO and stored in liquid nitrogen until staining for flow cytometry.

### Separation of lymphocytes from mouse spleen

After removing the capsular layer in a sterile petri dish, the spleen was mechanically disrupted using sterile BP blades. The cell suspension in RPMI1640 media (Corning, USA) was filtered through a cell strainer (70uM Nylon mesh, Fisherbrand, USA) to make a single-cell suspension. Cells were washed once in RPMI 1640 (Corning, USA) and then treated with ammonium chloride solution (STEMCELL Technologies, Canada) at a 9:1 ratio in RPMI 1640 cell culture media for 7 min on ice to lyse RBCs. After lysing the RBCs, the cells were washed twice with RPMI 1640 medium, resuspended in freezing media containing 10% DMSO, and stored in liquid nitrogen until further use.

### Flow cytometry analysis

On average, 10^5^ – 10^7^ cells were used for staining. A list of antibodies with fluorochrome was used, as in **(Supplementary. Table3)**. Stained cells were fixed in IC fixation buffer (Thermo Fisher, USA) and run on an Attune NxT Flow cytometer (Thermo Fisher, USA). Raw files were exported to the FlowJo software for further analysis.

### Histopathology sectioning and Scoring

Histological analysis was conducted on both the distal and proximal ends of the murine colon to evaluate the extent of tissue damage and the therapeutic effects of the interventions. Tissues were fixed in formalin, embedded in paraffin, and then sectioned at 10um for staining. Hematoxylin and Eosin (H&E) staining was performed to assess general morphology and inflammation, while Periodic Acid-Schiff (PAS) staining was used to highlight goblet cell abundance and mucin production. A board-certified pathologist, blinded to the experimental conditions, evaluated the stained sections.

The scoring system ranged from 0 (absent) to 4 (severe), based on the degree of inflammatory cell infiltration, epithelial damage, and mucosal architecture disruption observed. Scores of 1 indicated minimal changes, 2 mild changes, 3 moderate changes, and 4 severe pathologies. The scoring was independently performed for both the distal and proximal colon sections and reported based on inflammation.

## Statistical analysis

Statistical analyses were performed using GraphPad Prism 9 software (GraphPad Software, San Diego, CA, USA). Comparisons between groups were performed using either the Mann-Whitney t-test or t-test with Welch’s correction unless mentioned in the figure legends. Differences were considered statistically significant at a p value of 0.05. A bubble plot was plotted in R using the ggplot package.

## Data Availability Statement

All 16s amplicon sequencing data generated from this project were deposited in the NCBI SRA database under BioProject PRJNA1013066. Raw whole-genome sequence data for the strains used were obtained from a previously published study [27] under the BioProject <u>PRJNA494608</u>.

## Ethics Statement

All animal procedures were approved by the Institutional Animal Care and Use Committee of South Dakota State University with the Prior approval of protocol 19-014A.

## Author Contributions

AA, LA, AM, SG, and SM performed the experiments, VS and SM performed the pathological analysis, JS and PK designed the study and provided resources. AA wrote the manuscript with input from other authors.

## Acknowledgments

Computations supporting this project were performed on High-Performance Computing systems managed by Research Computing Group, part of the Division of Technology and Security at South Dakota State University.

## Supplementary Material

### Supplementary Figures

**Supplementary Figure 1.**
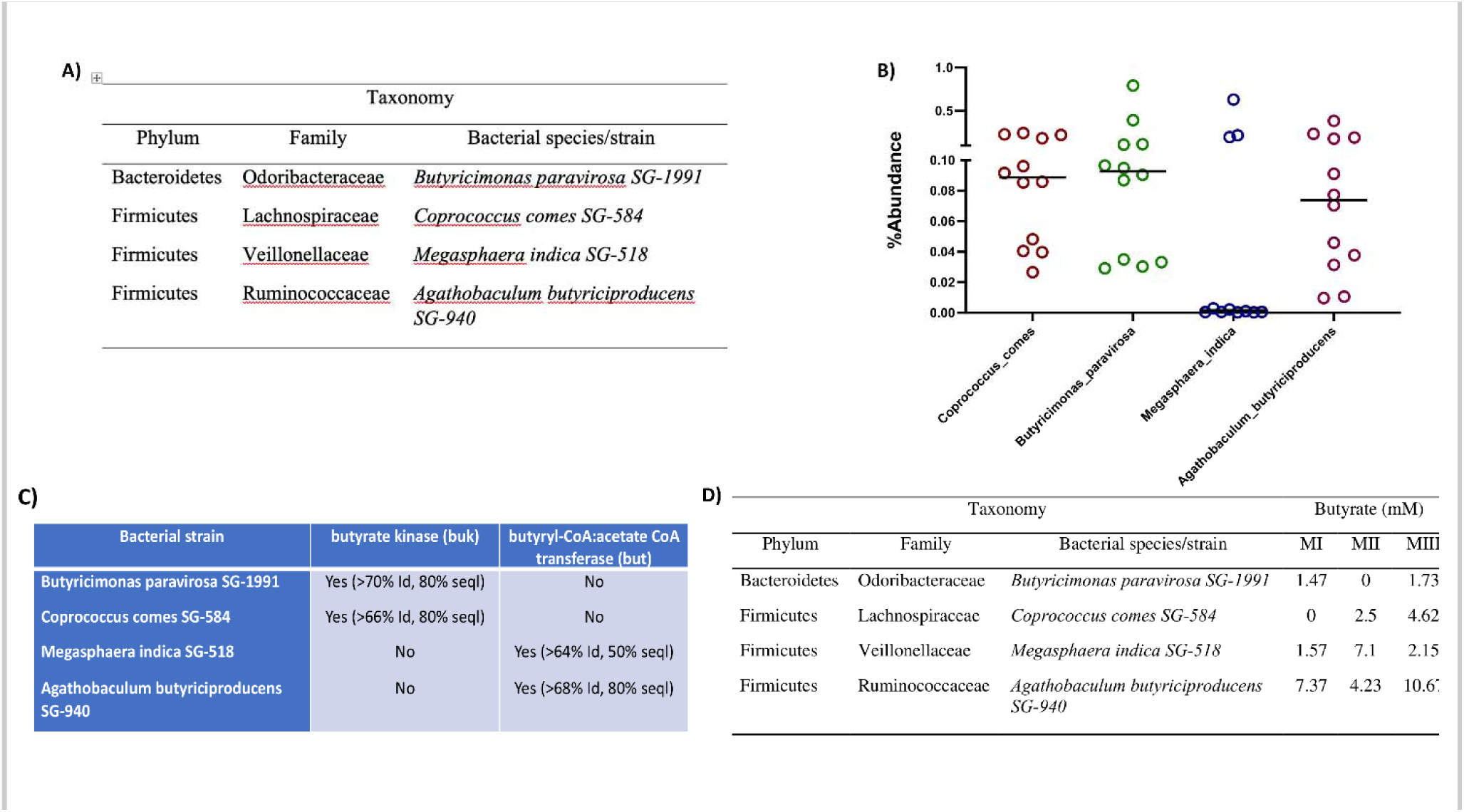
SC-4 species showed the ability to produce butyrate both genotypically and phenotypically. **(A)** Table showing the phylum, and family to which *Coprococcus comes*, *Agathobaculum butyriciproducens, Megasphaera indica,* and *Butyricimonas paravirosa* (SC-4) species belong. **(B)** Percentage abundance of SC-4 species in donor fecal sample from which it was isolated previously. **(C)** Table showing the presence/absence and percentage identity for buk and but genes in the genome of SC-4 species. **(D)** The concentration of butyrate produced by SC-4 strains invitro in different media conditions. (MI-mBHI supplemented with Inulin, MII-mBHI, MIII-DSMZ’s modified Peptone Yeast extract Glucose )

**Supplementary Figure 2.**
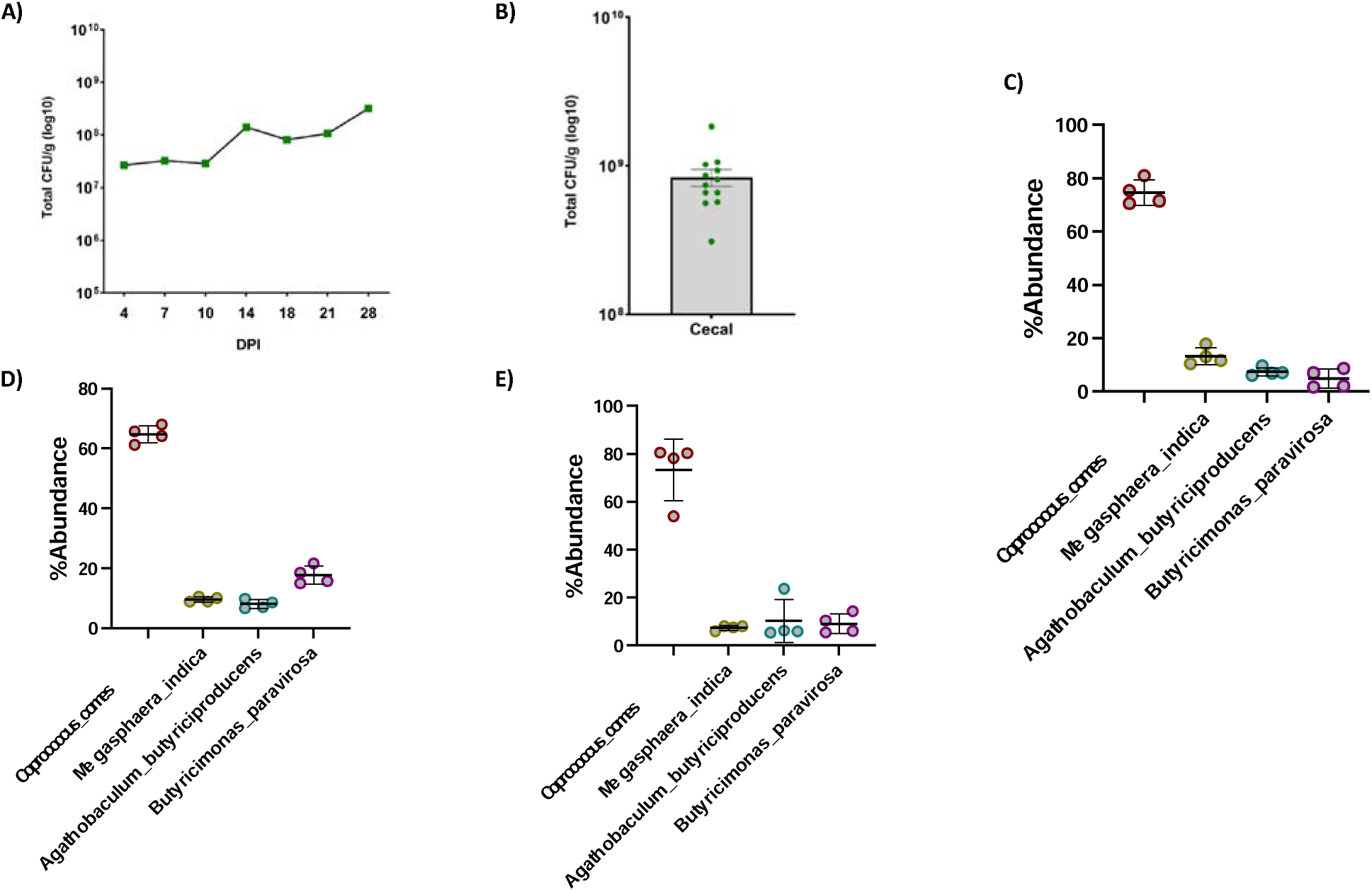
SC-4 successfully colonized GF mice. The total amount of bacteria found in **(A)** mice feces and Final Bacterial load in the **(B)** cecum post euthanasia (Day-28) was calculated by plating on PYG agar medium represented as Colony Forming Unit (CFU). Percentage abundance of *Coprococcus comes*, *Agathobaculum butyriciproducens, Megasphaera indica,* and *Butyricimonas paravirosa* (SC-4) in mice feces **(C)** 7 DPI, **(D)** 14 DPI, and **(E)** colon post euthanasia (Day-28).

**Supplementary Figure 3.**
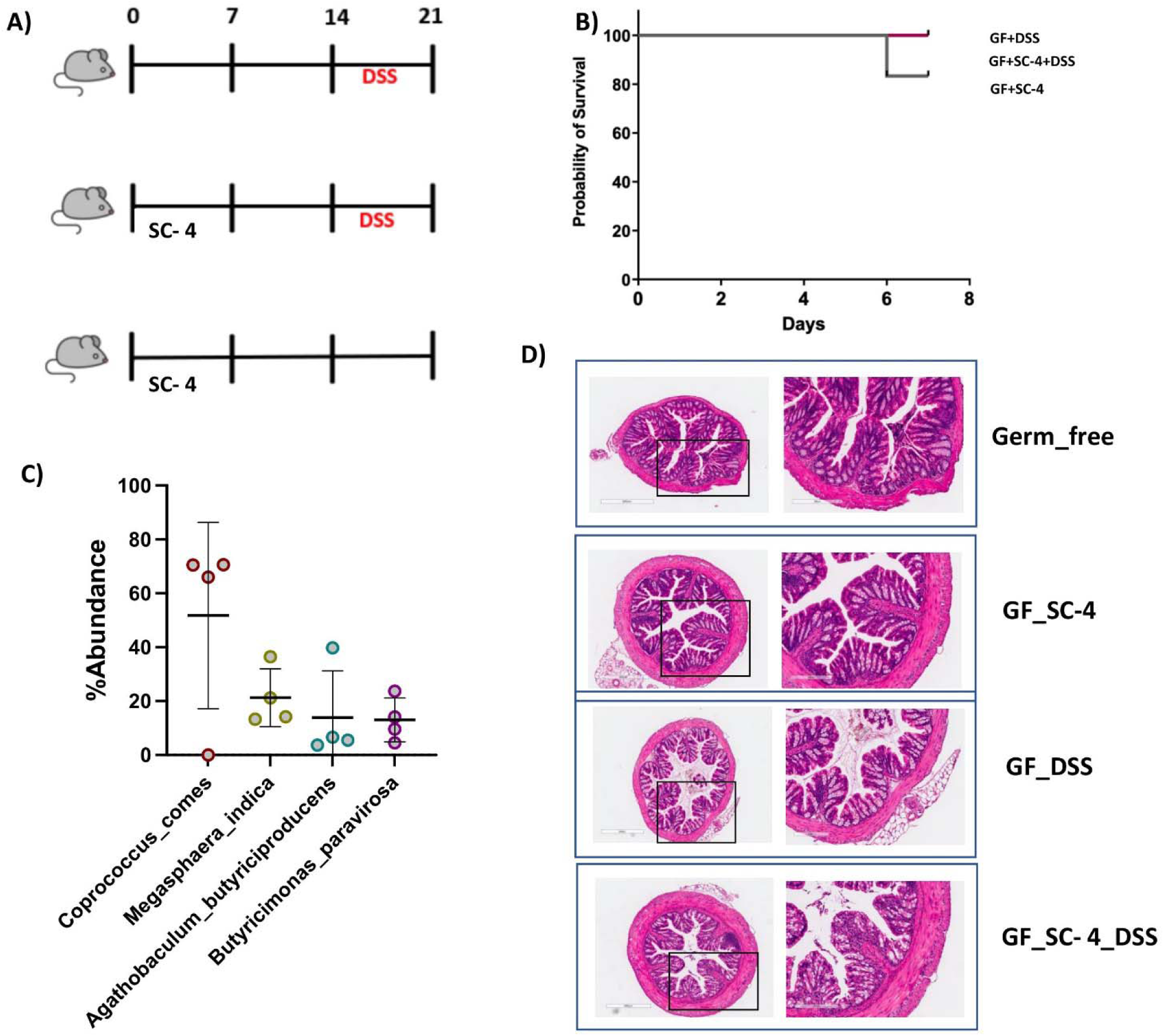
SC-4 protects against DSS-induced colitis. Experimental workflow for DSS-induced colitis model in **(A)** GF mice (see more details in methods). **(B)** Survival graph for GF mice induced with colitis (GF_DSS), SC-4 pre-colonized induced with colitis (GF+SC-4+DSS), or SC-4 pre-colonized only(GF+SC-4). **(C)** Percentage abundance of *Coprococcus comes*, *Agathobaculum butyriciproducens, Megasphaera indica,* and *Butyricimonas paravirosa* (SC-4) in mice colon post euthanasia (Day-21). Representative histopathological photograph showing the colon tissue cross-section after H&E staining of Germ free mice (GF) , GF mice induced with colitis (GF_DSS), GF mice gavaged with SC-4 bacteria (SF_SC-4) and SC-4 pretreated mice induced with colitis (GF_SC-4_DSS).

**Supplementary Figure 4.**
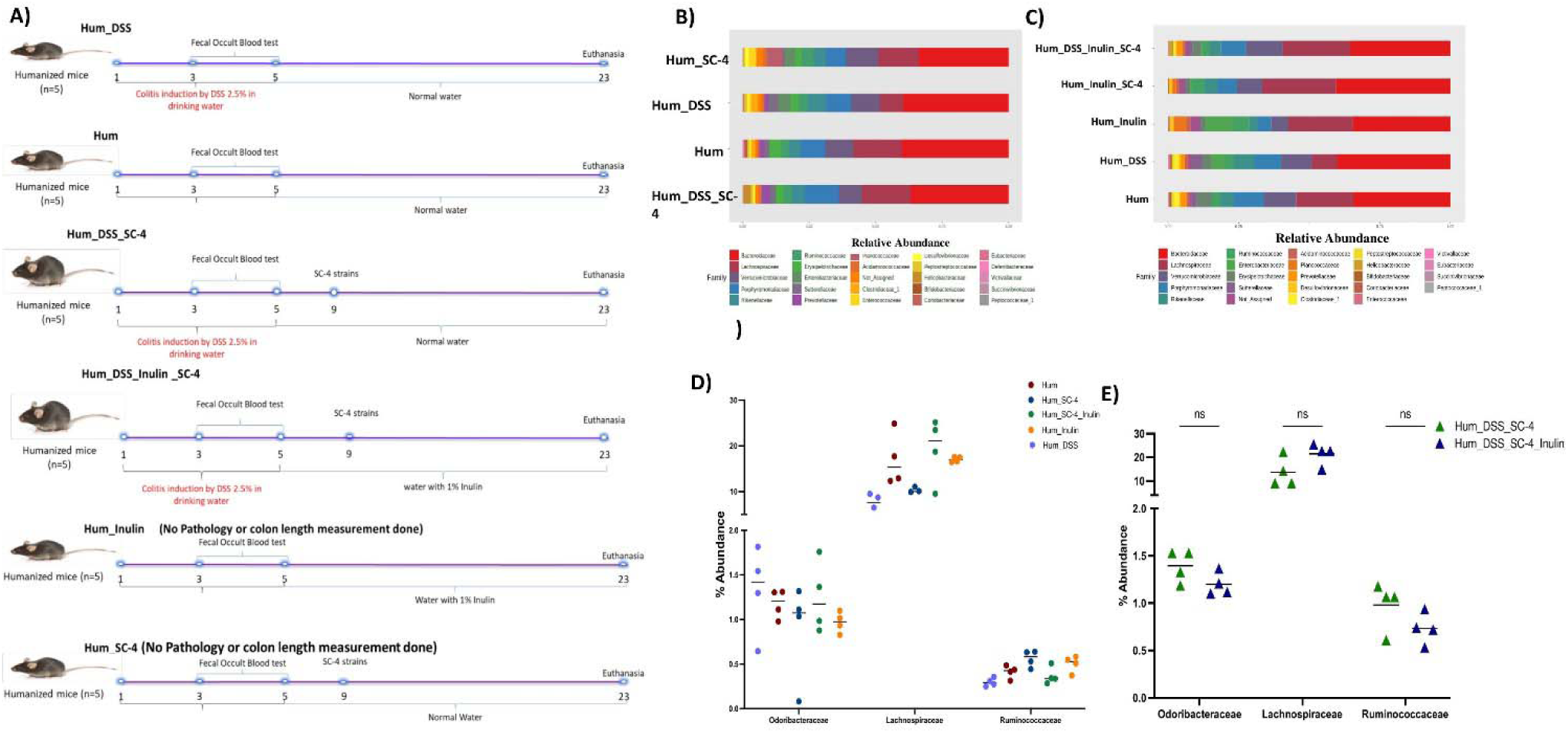
SC-4 help restore lost microbiota and help humanized mice recover from DSS-induced colitis. Experimental outline for humanized mice based mice experiment for humanized mice with DSS-induce colitis treated with SC-4 (Hum_DSS_SC-4), Humanized mice gavage with SC_4 (Hum_SC-4), humanized mice (Hum), humanized mice induced with colitis (Hum_DSS), humanized mice treated with SC-4 with supplementation of Inulin (Hum_Inulin_SC-4), humanized mice with DSS-induce colitis treated with SC-4 with supplementation of Inulin (Hum_DSS_Inulin_SC-4) and humanized mice with supplementation of Inulin (Hum_Inulin) **(A)**. Relative abundance of bacterial community in the cecum of mice for **(B)** Hum_SC-4, Hum_DSS, Hum or **(C)** Hum_DSS_SC-4 and Hum_DSS_Inulin_SC-4, Hum_Inulin_SC-4, Hum_Inulin, Hum_DSS, or Hum. Percentage abundance of *Butyricimonas paravirosa* (Odoribacteraceae), *Coprococcus comes* (Lachnospiraceae), or *Agathobaculum butyriciproducens* (Ruminococcaceae) in mice cecum for Hum, Hum_SC-4, Hum_SC-4_Inulin, Hum_Inulin, or Hum_DSS **(D)** and Hum_DSS_SC-4 and Hum_DSS_SC-4_Inulin groups post Euthanasia.

**Supplementary Figure 5.**
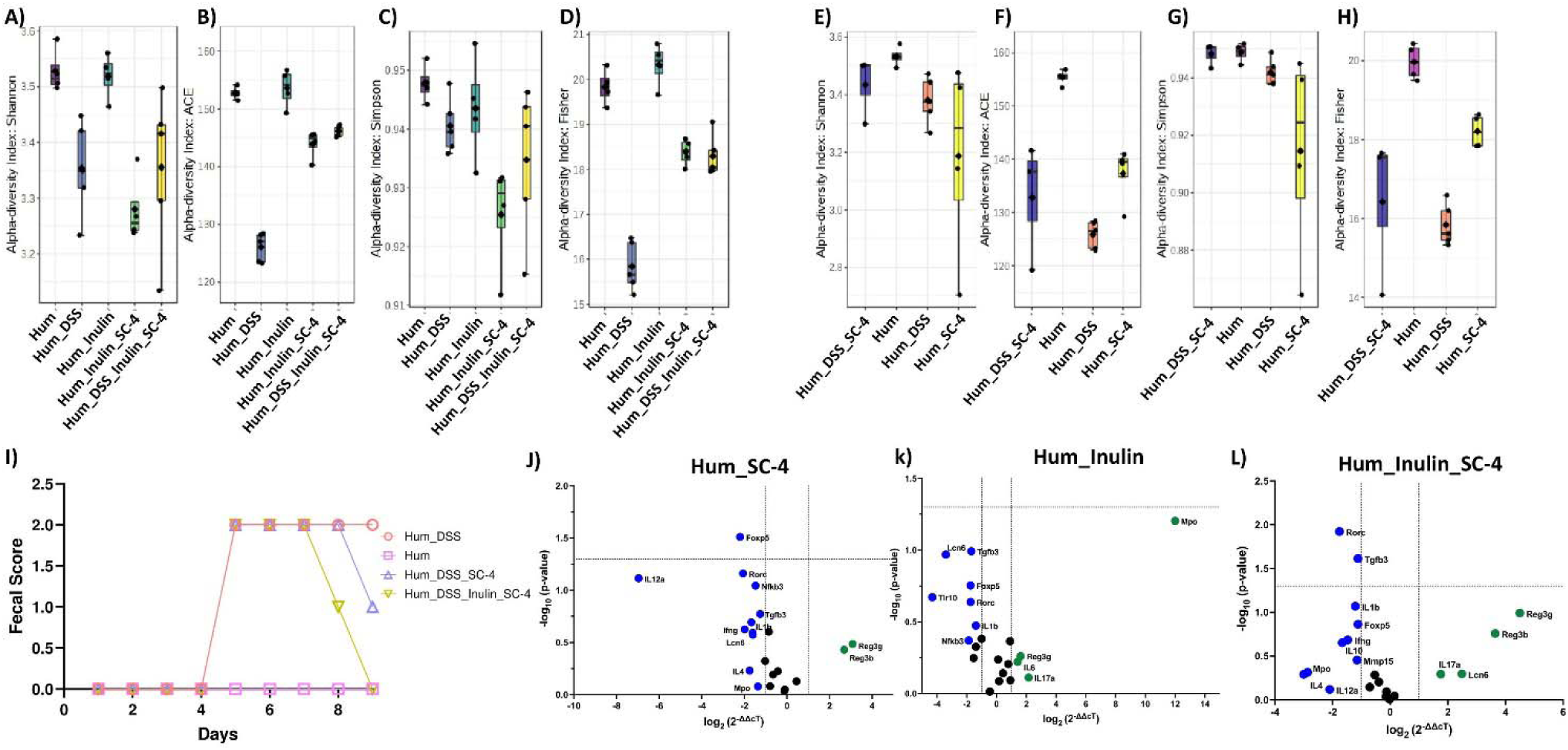
SC-4 increased the diversity of the gut microbiome in humanized mice. Alpha diversity analysis (Shannon, ACE, Simpson, or Fisher) for humanized mice treated with SC-4 with supplementation of Inulin (Hum_Inulin_SC-4), humanized mice with DSS-induce colitis treated with SC-4 with supplementation of Inulin (Hum_DSS_Inulin_SC-4) and humanized mice with supplementation of Inulin (Hum_Inulin) **(A-D),** humanized mice with DSS-induce colitis treated with SC-4 (Hum_DSS_SC-4), Humanized mice gavage with SC_4 (Hum_SC-4), humanized mice (Hum), humanized mice induced with colitis (Hum_DSS) **(E-H)** Fecal Score recorded for Hum_DSS, Hum, Hum_DSS_SC-4, and Hum_DSS_Inulin_SC-4 groups post DSS treatment. RT-PCR gene expression for selected 25 immune-related genes of mice colon for **(J)** Hum_SC-4, **(K)** Hum_Inulin, and **(L)** Hum_Inulin_SC-4. Green dots - overexpressed more than 2-fold change, Blue dots-represses more than 2-fold compared to Hum control (dotted lines on the x-axis represent a 2-fold increase or decrease in expression and the y-axis represents a p-value cutoff of 0.05).

**Supplementary Figure 6.**
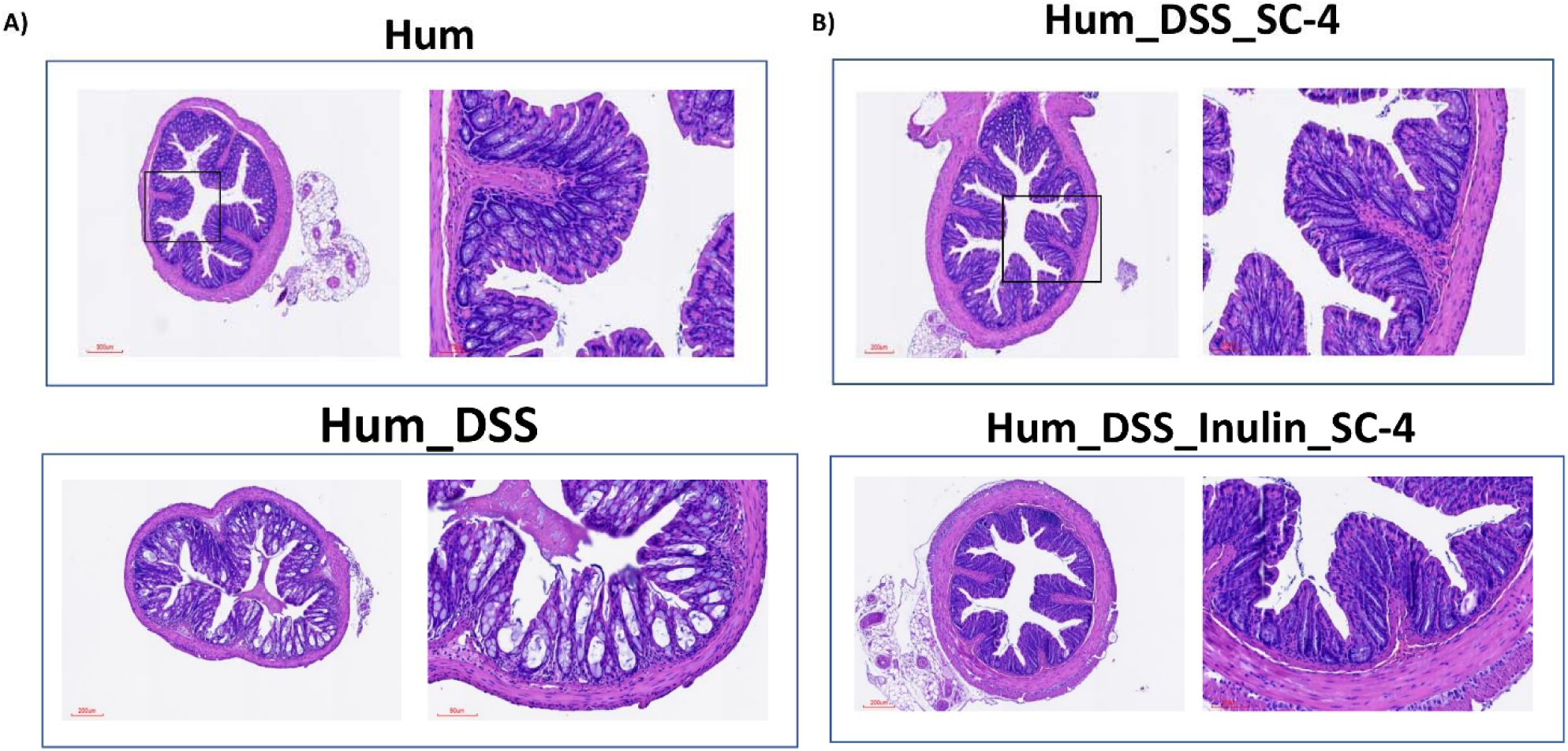
SC-4 treatment help recover colon tissue damage caused by colitis. Representative histopathological photograph showing the colon tissue cross-section after H&E staining for humanized mice (Hum), humanized mice induced with colitis (Hum_DSS) **(A)** humanized mice with DSS-induce colitis treated with SC-4 (Hum_DSS_SC-4) and humanized mice with DSS-induce colitis treated with SC-4 with supplementation of Inulin (Hum_DSS_Inulin_SC-4) and humanized mice with supplementation of Inulin (Hum_Inulin) **(C,E)**. Representative colon images from Hum, Hum_DSS, Hum_DSS_SC-4, and Hum_DSS_Inulin_SC_4 **(F)** and associated Violin plot **(G)** representing the length of the colon in cm. Statistical significance for (G) calculated using one-way annova with Dunnet correction with absolute p-value represented on the graph. Bubble plot representing relative abundance in natural log scale at a species level resolution for SCFA producing bacteria (Similar to Figure 1B) from Hum, Hum_DSS, Hum_DSS_SC-4, and Hum_DSS_Inulin_SC_4 mice groups **(H)**.

## Supplementary Tables

**Supplementary Table 1.**
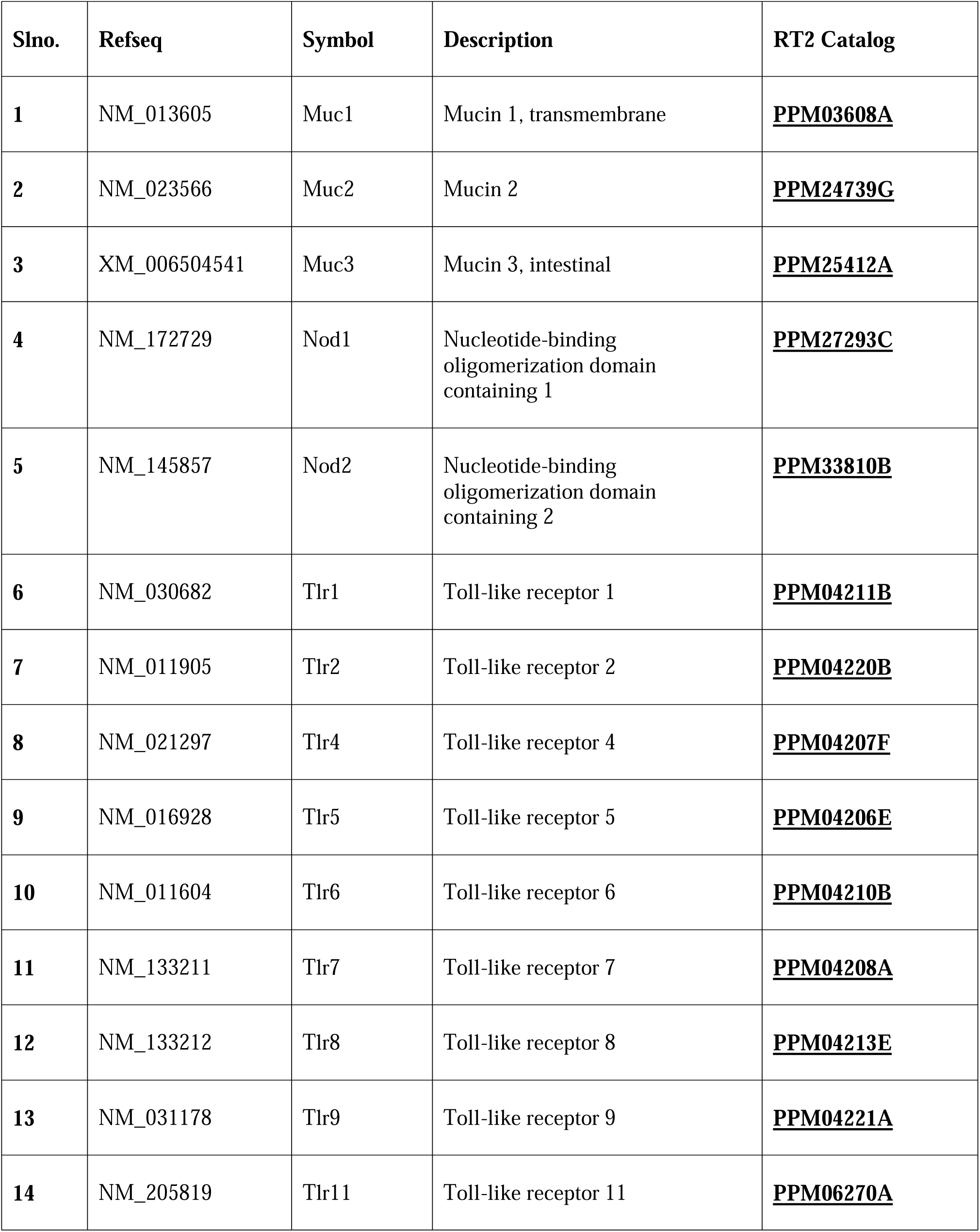
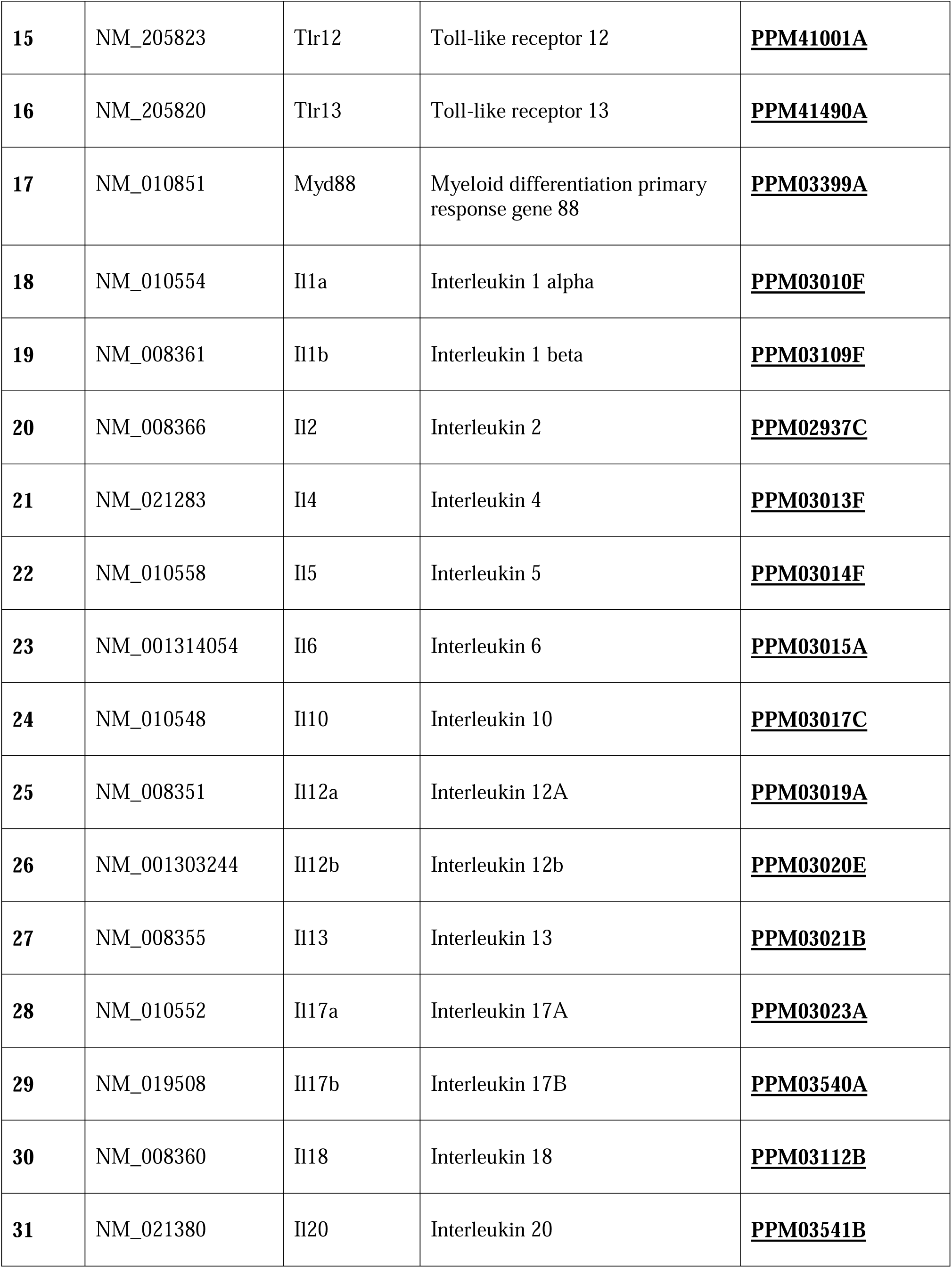
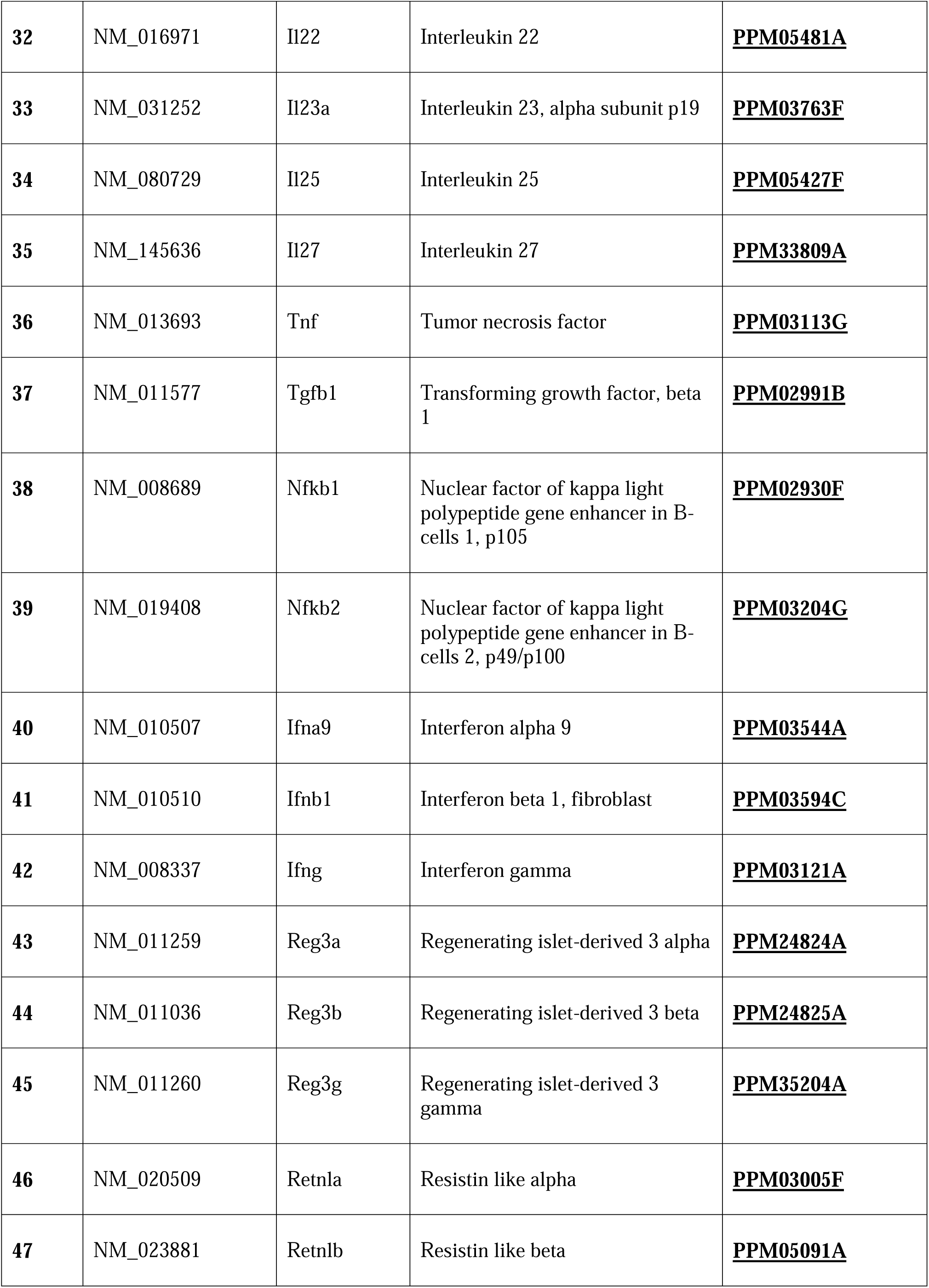
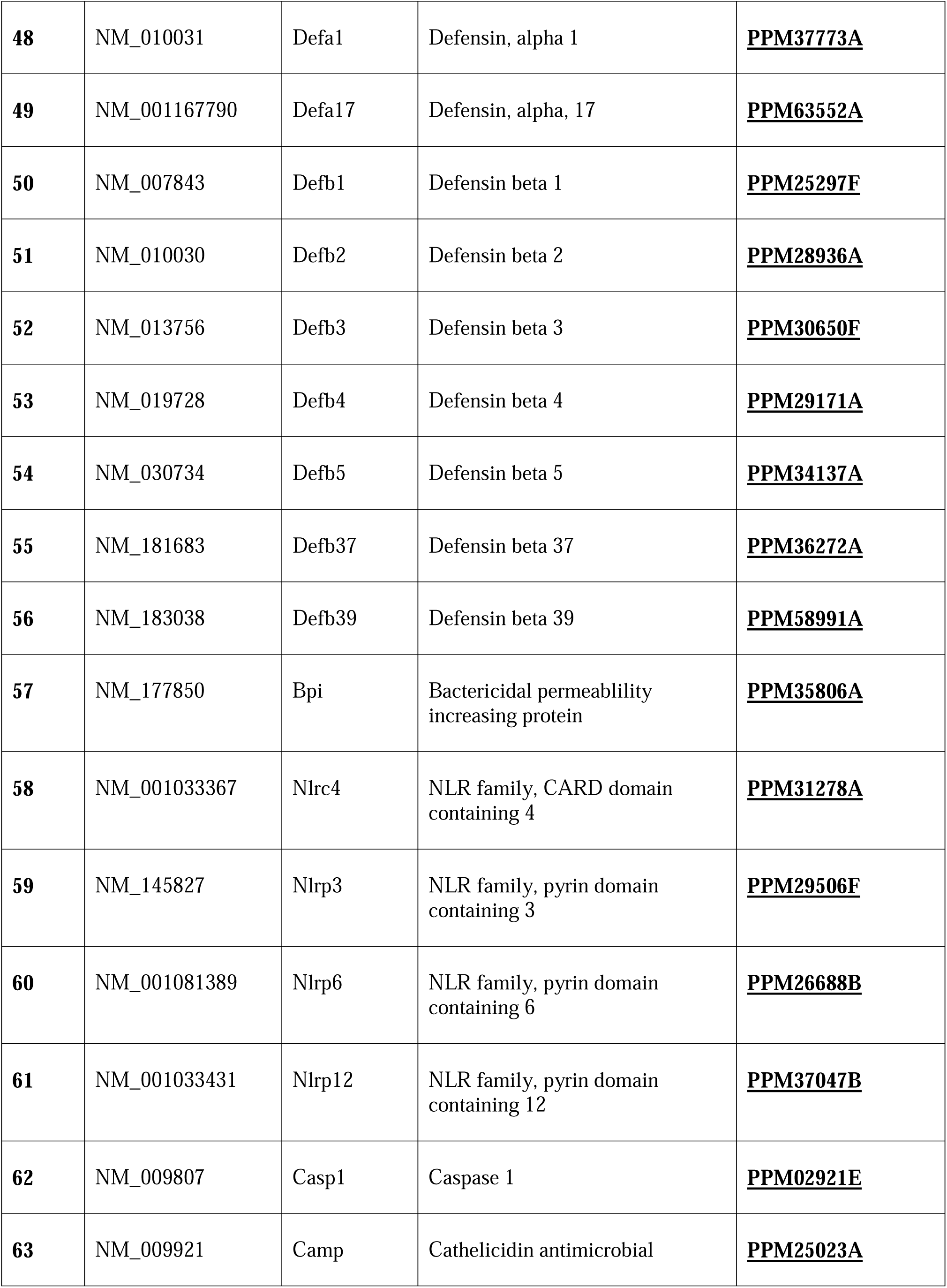
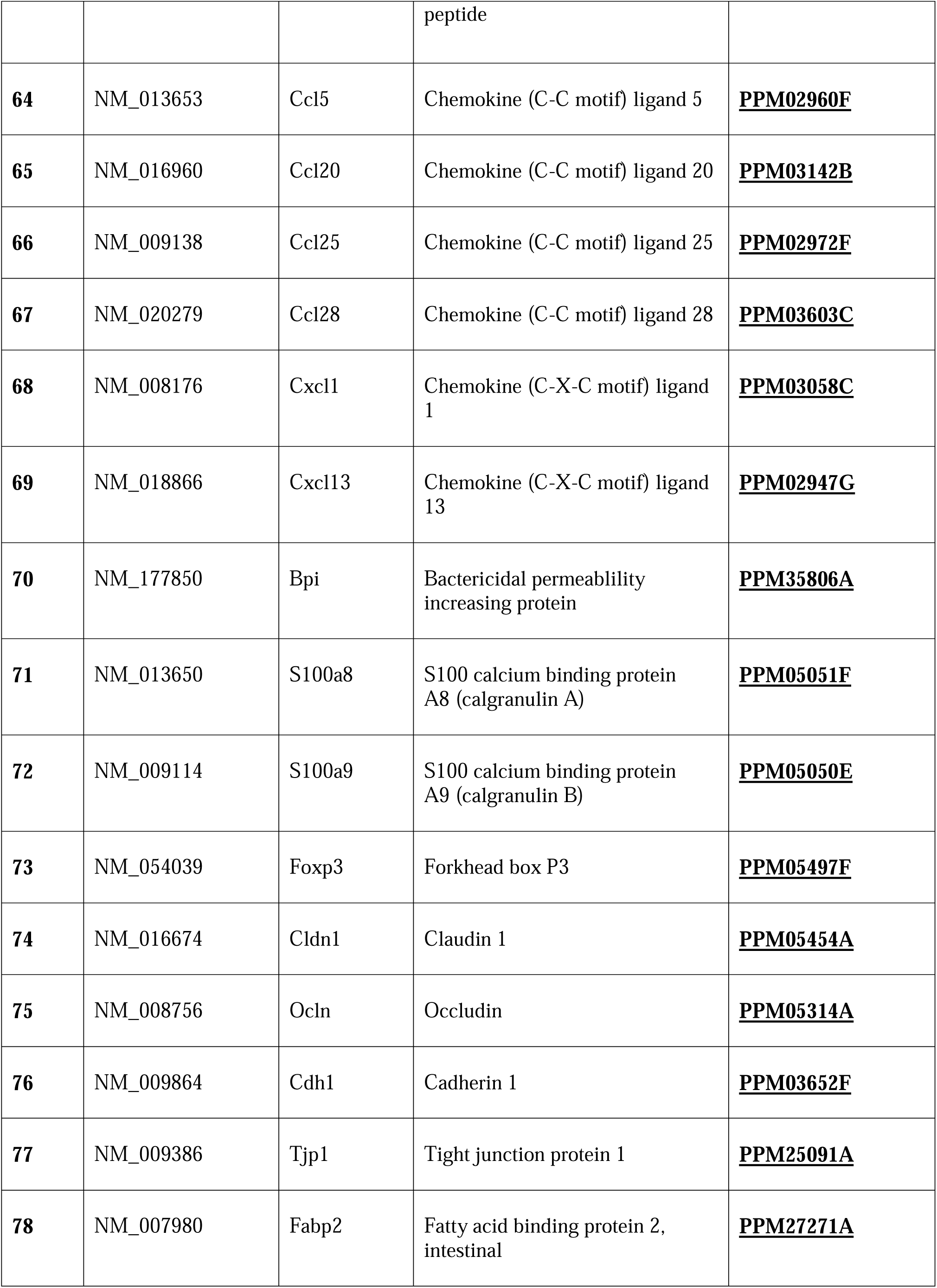
Gene names used in Rt2 profiler array gene expression analysis.

**Supplementary Table 2.**
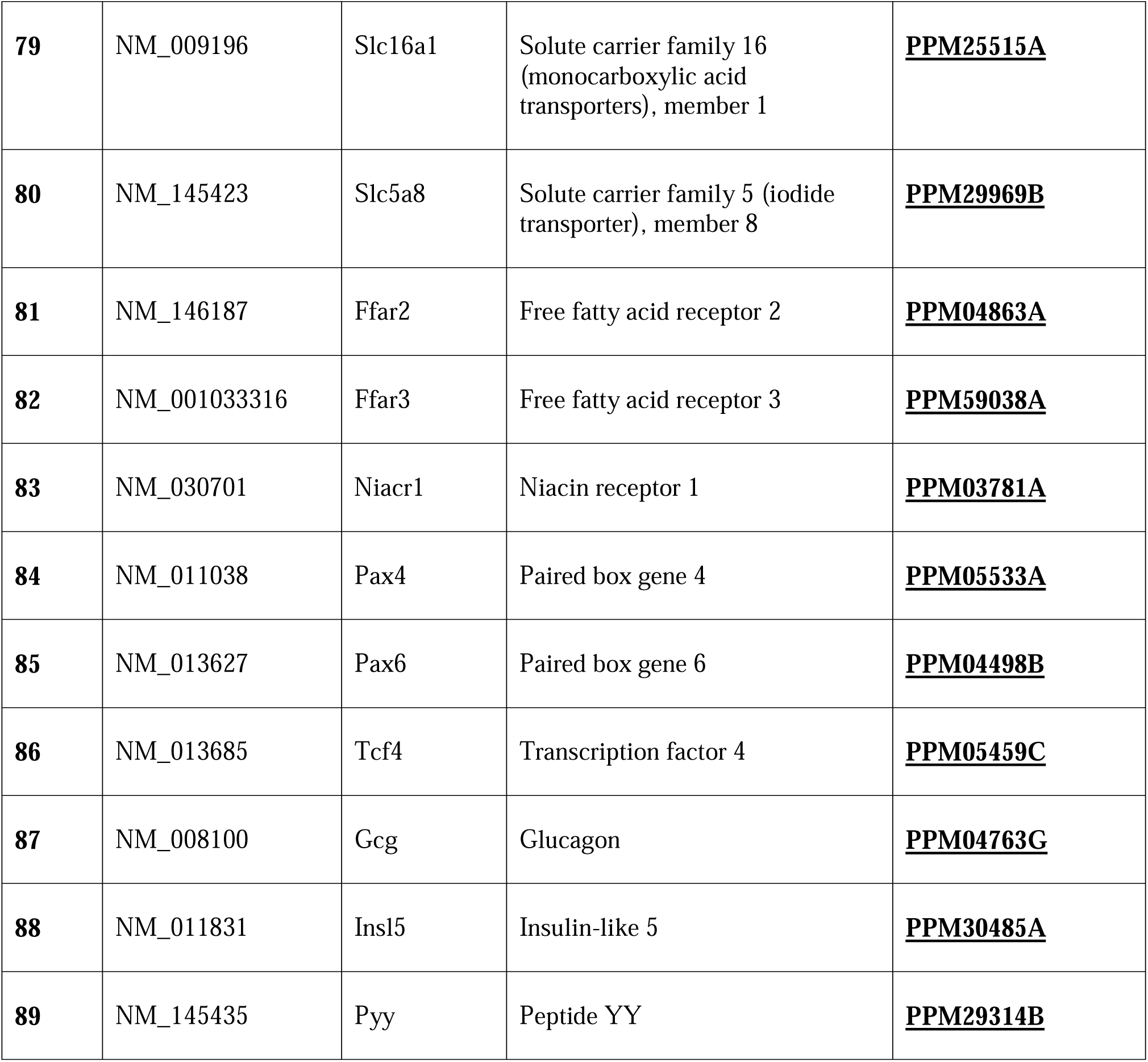
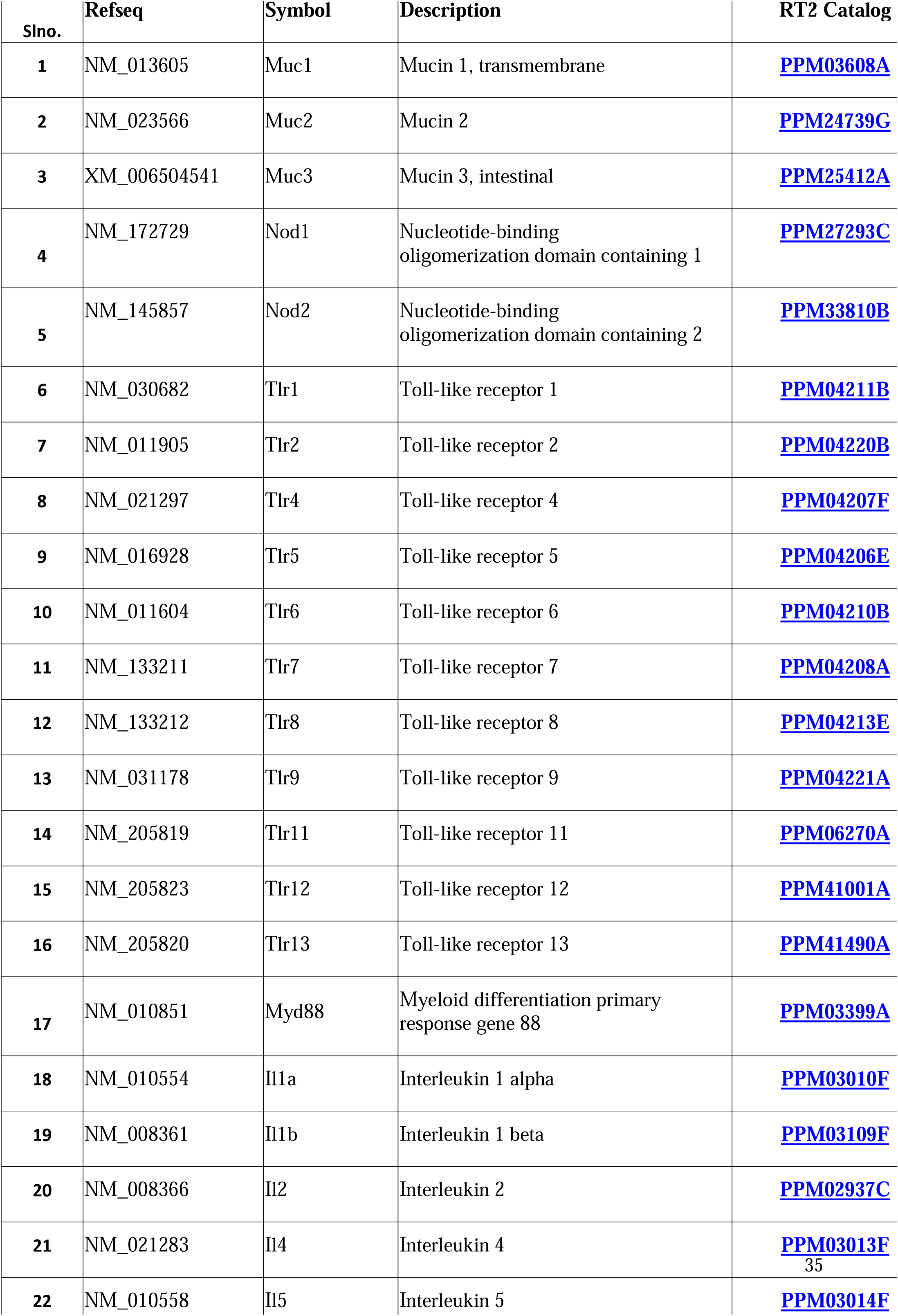
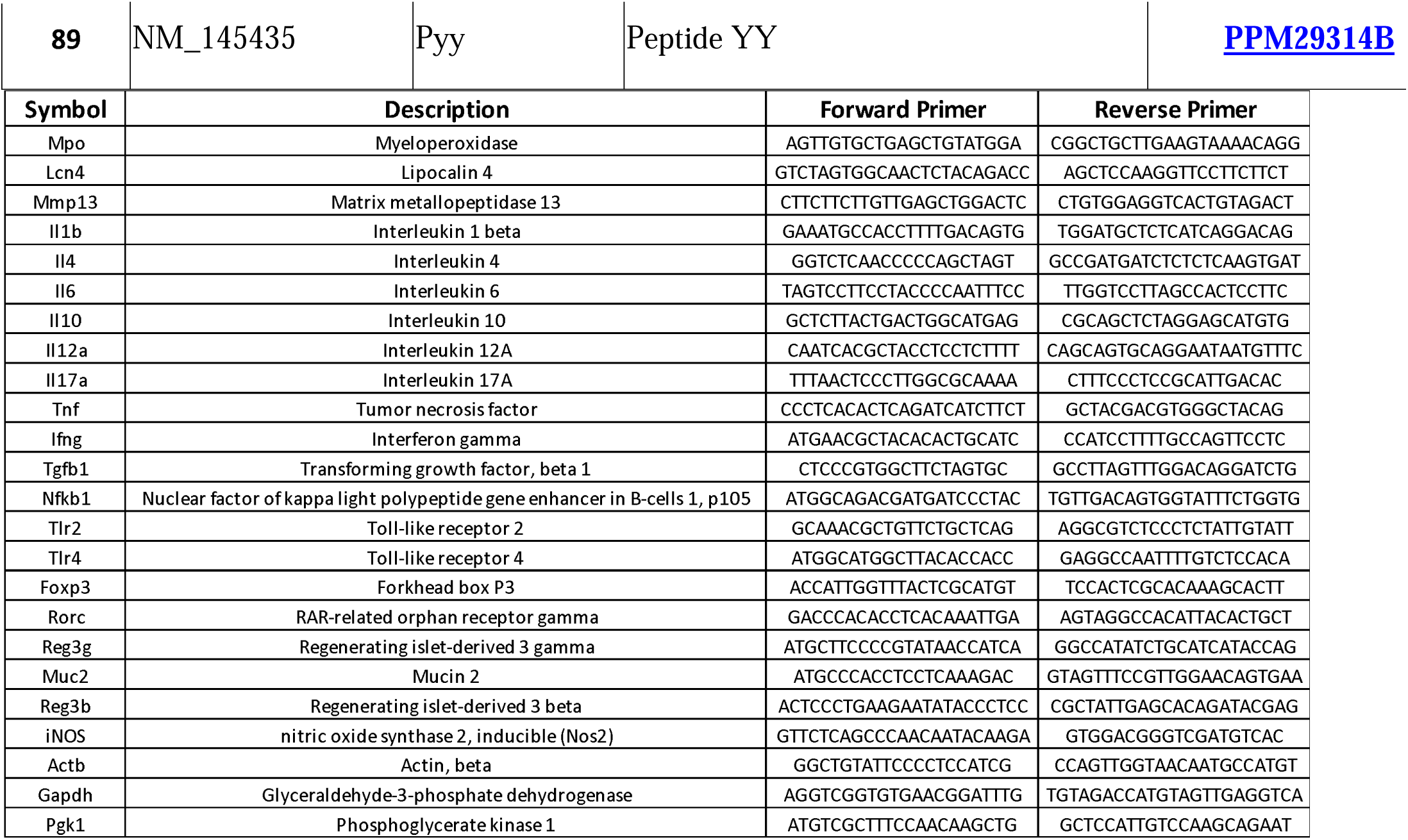
Primer details of the custom-made panel of mice immune genes.

**Supplementary Table 3.** List of antibodies with fluorochrome used for staining the immune cells under panel 1&2.

| Fluorochrome | Cell type | Antigen | Clone | Product |
| --- | --- | --- | --- | --- |
| **Panel -1** |  |  |  |  |
| FITC | Lymphocytes total | CD45 | 30-F11 | CD45 Monoclonal Antibody (30-F11), FITC, eBioscience |
| PE | B cells | CD19 | eBio1D3 (1D3) | CD19 Monoclonal Antibody (eBio1D3(ID3)), PE, eBioscience |
| PE-Cyanine7 | Tgd | TCR gamma/delta | eBioGL3 (GL-3,GL3) | TCR gamma/delta Monoclonal Antibody (eBioGL3(GL-3,GL3)), PE-Cyanine7, eBioscience |
| APC | ILC3 | ROR gamma (t) | AFKJS-9 | ROR gamma (t) Monoclonal Antibody (AFKJS-9), APC, eBioscience |
| Alexa Fluor 700 | Tab | TCR beta | H57-597 | TCR beta Monoclonal Antibody (H57-597), Alexa Fluor 700, ebioscience |
| LIVE/DEAD Fixable Near-IR Dead Cell Stain | Live cells |  |  | LIVE/DEAD Fixable Near-IR Dead cell Stain Kit, for 633 or 635 nm excitation |
| **Panel -2** |  |  |  |  |
| FITC | Lymphocytes total | CD45 | 30-F11 | CD45 Monoclonal Antibody (30-F11), FITC, eBioscience |
| PE | CD8 cells | CD8 | 53-6.7 | CD19 Monoclonal Antibody (53-6.7), PE, eBioscience |
| PE-Cyanine5.5 | CD4 cells | CD45 | RM4-5 | CD4 Monoclonal Antibody (RM4-5),PE-Cyanine5.5, eBioscience |
| PE-Cyanine7 | Treg | FOXP-3 | FJK-16s | FOXP3 Monoclonal Antibody (FJK-16s),PE-Cyanine7, eBioscience |
| APC | Th17 | ROR gamma (t) | AFKJS-9 | ROR gamma (t) Monoclonal Antibody (AFKJS-9), APC, eBioscience |
| Alexa Fluor 700 | Tab | TCR beta | H57-597 | TCR beta Monoclonal Antibody (H57-597), Alexa Fluor 700, ebioscience |
| LIVE/DEAD Fixable Near-IR Dead Cell Stain | Live cells |  |  | LIVE/DEAD Fixable Near-IR Dead cell Stain Kit, for 633 or 635 nm excitation |

